# AP-1 Mediates Cellular Adaptation and Memory Formation During Therapy Resistance

**DOI:** 10.1101/2024.07.25.604999

**Authors:** Jingxin Li, Pavithran T. Ravindran, Aoife O’Farrell, Gianna T. Busch, Ryan H. Boe, Zijian Niu, Sean Woo, Margaret C. Dunagin, Naveen Jain, Yogesh Goyal, Kavitha Sarma, Meenhard Herlyn, Arjun Raj

**Affiliations:** Genetics and Epigenetics, Cell and Molecular Biology Graduate Group, Perelman School of Medicine, University of Pennsylvania, Philadelphia, PA, USA; Cancer Biology Program, Cell and Molecular Biology Graduate Group, Perelman School of Medicine, University of Pennsylvania, Philadelphia, PA; Department of Bioengineering, School of Engineering and Applied Sciences, University of Pennsylvania, Philadelphia, PA, USA; Department of Chemistry, School of Arts and Sciences, University of Pennsylvania, Philadelphia, PA, USA; Department of Physics and Astronomy, School of Arts and Sciences, University of Pennsylvania, Philadelphia, PA, USA; Computational and Systems Biology Program, Massachusetts Institute of Technology, Cambridge, MA, USA; Department of Biology, School of Arts and Sciences, University of Pennsylvania, Philadelphia, PA; Department of Genetics, Perelman School of Medicine, University of Pennsylvania, Philadelphia, PA, USA; Department of Cell and Developmental Biology, Feinberg School of Medicine, Northwestern University, Chicago, IL, USA; Center for Synthetic Biology, Northwestern University, Chicago, IL, USA; Robert H. Lurie Comprehensive Cancer Center, Northwestern University Feinberg School of Medicine, Chicago, IL, USA; The Wistar Institute, Gene Expression and Regulation program, Philadelphia, PA, USA; Department of Cell and Developmental Biology, Perelman School of Medicine, University of Pennsylvania, Philadelphia, PA, USA; The Wistar Institute, Molecular and Cellular Oncogenesis Program and Melanoma Research Center, Philadelphia, PA, USA

## Abstract

Cellular responses to environmental stimuli are typically thought to be governed by genetically encoded programs. We demonstrate that melanoma cells can form and maintain cellular memories during the acquisition of therapy resistance that exhibit characteristics of cellular learning and are dependent on the transcription factor AP-1. We show that cells exposed to a low dose of therapy adapt to become resistant to a high dose, demonstrating that resistance was not purely selective. The application of therapy itself results in the encoding of transient gene expression into cellular memory and that this encoding occurs for both transiently induced and probabilistically arising expression. Chromatin accessibility showed concomitant persistence. A two-color AP-1 reporter system showed that these memories are encoded in *cis*, constituting an example of activating *cis* epigenetics. Our findings establish the formation and maintenance of cellular memories as a critical aspect of gene regulation during the development of therapy resistance.

## Introduction

Cells are thought to regulate their function by executing prescribed programs that are hard-coded in their DNA through the interactions of signaling and gene regulatory proteins. A classic example is the regulation of the Lac operon in response to lactose^1,2^, presumably shaped by millions of years of evolution to mediate the proper cellular responses to various stimuli. Although these programs can be quite intricate, they are typically immutable in that the programs themselves do not change, akin to instinctual behaviors. In contrast, other biological control systems, perhaps most notably networks of neurons, have not only instinctual but also learned behaviors that allow flexible responses to new situations that may never have been encountered before. Cellular learning within single cells, outside of a few contexts^3–5^ (most notably inflammatory responses^6,7^), has remained controversial.

In the cellular context, we define memory as an encoding of a previous state of the cell and learning as reading out (and potentially acting upon) that memory. There are two broad categories of memory that can be encoded and retained, which, in the language of epigenetics, correspond to the distinction between “*trans*” and “*cis*” epigenetics^8^. *Trans* epigenetics refers to memories that are encoded in the overall state of the cell exogenous to the genome itself and could be equated to the execution of a regulatory program. For example, a gene’s expression could be “remembered” through positive feedback if it encoded a transcription factor that enhanced its own transcription. These forms of memory are “hard-coded” into the DNA in the sense that if the DNA sequence were changed (say, of the transcription factor or its binding site), then the positive feedback could be lost. *Cis* epigenetics, on the other hand, refers to regulation by factors that are, to some extent, independent of the DNA sequence *per se*. Examples of *cis* epigenetics include persistent changes in expression mediated by self-propagating non-sequence-related changes to the gene itself, such as DNA methylation or some forms of histone modification, such as silencing of the X chromosome in females by Xist^9^ and genomic imprinting^10^.

Instances of *cis* epigenetics generally comprise examples of silencing, although there are examples of *cis* epigenetic propagation of activation, most notably in yeast^11–14^ and in some instances in flies^15^. In principle, *cis* epigenetics can flexibly encode memories that are not hard-coded into the sequence, with the specificity of memory formation relying on information beyond the sequence. However, it remains unclear whether cells exploit such flexibility, which is critical to learning, to perform functions beyond what is already specified in a set developmental or signal-response trajectory.

Here, we develop the concept of cellular learning in the context of therapy resistance in cancer cells. Typically, therapy resistance is thought to be driven by genetic differences, which would suggest a change to the hard-coded regulatory framework of the cell. Work from our lab and others, however, has shown that therapy resistance can arise from rare “primed” cells with non-genetic differences^16–26^. The existence of non-genetic differences suggests different possibilities for the acquisition of therapy resistance: cells primed for resistance could simply be selected by the therapy (as per natural selection of existing variation), or cells primed for resistance could change in some adaptive fashion, ultimately arriving at a fully therapy-resistant fate that is different from the initial primed state. Our prior results suggest the latter scenario, with both molecular and phenotypic differences between primed and fully resistant cells. Although this is consistent with other recent results^27^, a definitive demonstration of cellular adaptation has proven challenging.

Cellular adaptation itself could occur via two conceptually distinct modes corresponding to *trans* and *cis* epigenetics. In the first, cellular adaptation is the result of the execution of an inbuilt therapy resistance program, a cascade of hard-coded interactions that lead to a stably resistant cell state. The other possibility, however, is that cellular adaptation is the result of acting upon the encoding of memories of the state of the cell at the time of drug application. Our suspicion that cellular learning may underlie the process arose from the fact that these cancer cells do not have any natural history of exposure to targeted therapies, suggesting that they would be unlikely to possess any hard-coded program to specifically handle these insults. However, there is no direct evidence for cellular learning operating in this context.

Here, we demonstrate that in the process of becoming therapy-resistant, cells display various characteristics of cellular memory formation and learning. They show an adaptive phenotypic behavior, exhibiting the ability to resist higher doses of therapy than pure selection would predict. They also show the ability to encode the memory of the transcriptional state of the cell at the time of therapy application, maintaining and even enhancing the expression of otherwise transiently expressed genes (both induced and stochastically expressed). We demonstrate that the transcription factor AP-1 is an important mediator of this memory effect. Further, we show that AP-1 reporter genes encode memories in *cis*, meaning that the memory is encoded based on the instantaneous activity of the gene and not the DNA sequence alone. Together, our results establish the formation and maintenance of cellular memories as an important aspect of gene regulation in therapy resistance and suggest molecular mechanisms that may be responsible for cellular memory and learning in other biological contexts as well.

## Results

### Tracking profiles of cells through therapy resistance suggests an adaptive process

To probe the potential for cellular learning, we used a cellular model for non-genetic resistance to cancer therapy. Specifically, we used a melanoma cell line WM989^28^ in which the vast majority of cells die upon MAPK pathway inhibition (using either the BRAF^V600E^ inhibitors vemurafenib/dabrafenib or the MEK inhibitor trametinib). Despite having clonally bottlenecked the cell line, however, a small fraction (around 1 in 2000) of the cells (which we dub “primed” for resistance) survive the therapy treatment and resume proliferation after a short period of quiescence, ultimately forming a therapy-resistant colony in cell culture. We have assembled multiple lines of evidence that there is no genetic basis for these rare cells becoming therapy-resistant, including Luria-Delbrück fluctuation analysis^16^ and DNA sequencing (both targeted panel^16^ and whole genome sequencing^20^ of multiple therapy-naive and therapy-resistant clones). Instead, we have demonstrated that the rare-cell expression of certain sets of genes is strongly associated with cells being primed for therapy resistance^16,18^. The therapy induces a strong selection for survival or death based on the state of the cell (primed or not primed, respectively) before the therapy is initiated, but the primed cells could follow different trajectories as therapy treatment continues. One possibility is that the primed cell is selected by the therapy and expands but does not change; i.e., the primed cell is the same as the resistant cell. Such a scenario is typically the case in instances of selection of genetic mutants. However, another possibility is that primed cells undergo a transformation during their time in therapy akin to cellular adaptation, such that the fully therapy-resistant cell is different from the initial primed cell (Fig. 1A).

**Figure 1:**
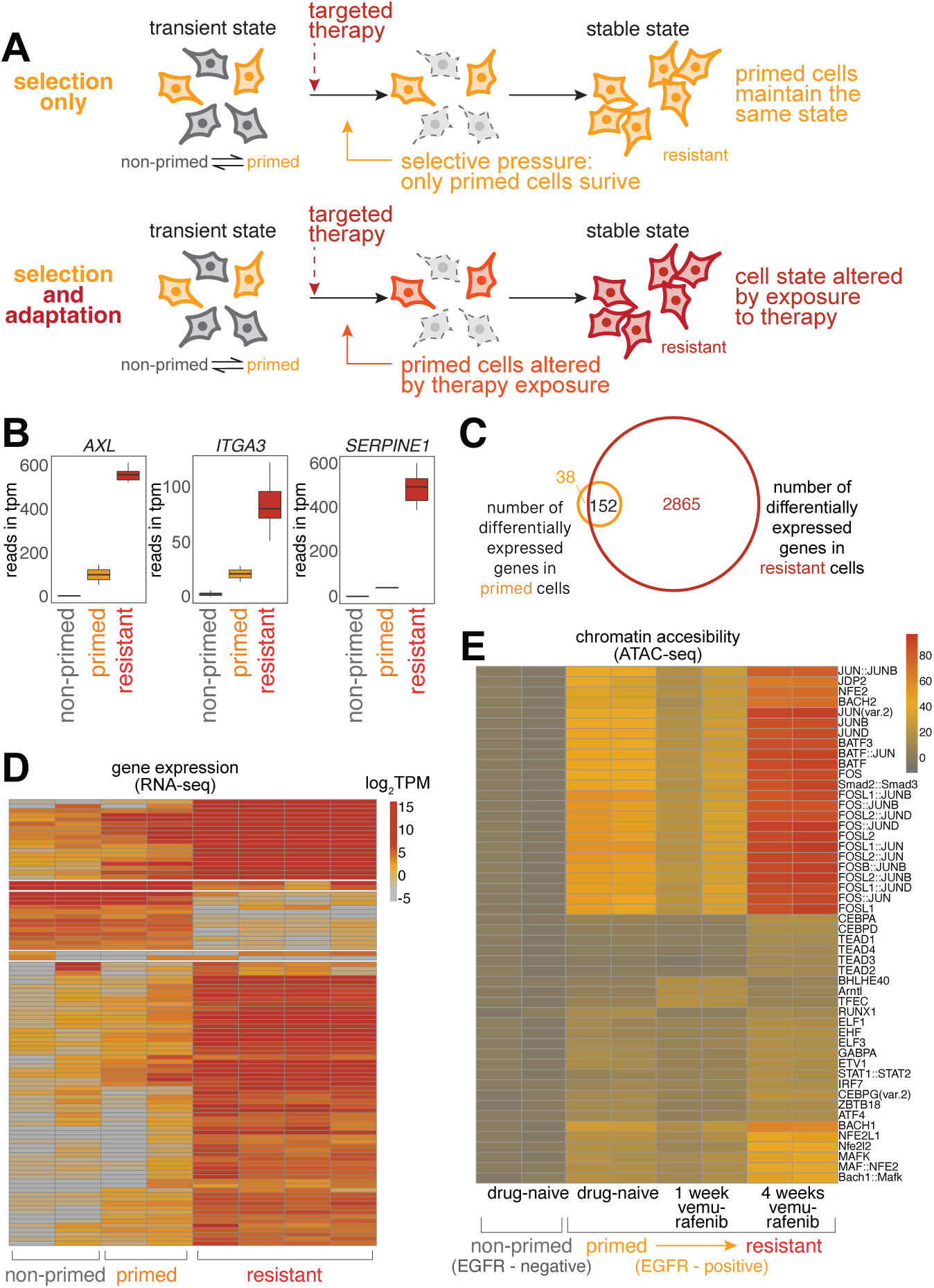
Adaptation and learning in WM989 melanoma cells. **(A)** Schematic comparison of resistance mechanisms in WM989 melanoma cells. Upper panel: Pure selection process where primed cells survive targeted therapy without altering their state. Lower panel: Adaptive learning process in which primed cells modify their cellular state in response to treatment, resulting in a molecularly and phenotypically distinct resistant population. **(B)** Transcriptional differences between non-primed, primed, and fully resistant cells measured by RNA-seq. Selected examples include previously identified priming marker genes. Resistant cells are those remaining after treatment with 1 μM vemurafenib for 21 days. Primed cells refer to cells identified as twins of those in the naive population that will eventually become resistant, with RNA FISH probes targeting barcodes identified from the resistant cell population. Non-primed cells refer to the rest of the therapy-naive population; in this case, those without matching resistant cell barcodes. **(C)** Venn diagram showing 3,017 differentially expressed genes in resistant cells (as compared to therapy-naive, non-primed cells) and 190 differentially expressed genes in therapy-naive primed cells. 152 of the 190 differentially expressed genes in primed cells are also present in the set of differentially expressed genes in resistant cells (c.f. Emert et al., 2021)^18^. **(D)** Heatmap of top variable genes from RNA-seq analysis in the global analysis of non-primed, primed, and fully resistant cells (as defined in panel B). **(E)** Heatmap of chromVAR analysis of ATAC-seq data. Primed cells in this experiment were identified by the expression of the surrogate marker EGFR. Top 0.02–0.2% of live cells staining for EGFR were designated as EGFR-high. (left to right) Two biological replicates for EGFR-negative (non-primed) cells never exposed to the therapy, EGFR-high (primed) cells never exposed to the therapy, EGFR-high (primed) cells exposed to 7 days of 1 μM vemurafenib, and EGFR-high (primed) cells exposed to 28 days of 1 μM vemurafenib.

A number of lines of evidence suggest the latter scenario. In our previous work, at the phenomenological level, we have shown that primed cells are only transiently primed^16,17,29^ while fully resistant cells remain resistant for months in continuous culture^16,20^. This demonstrated that cells change their phenotype between being primed and becoming fully resistant. To see how such changes manifested at the molecular level, we reanalyzed the transcriptomes of non-primed, primed, and fully resistant cells by retrospective identification of primed cells^18^ (Fig. 1B-D). As in Shaffer et al. (2017)^16,18^ and Emert et al. (2021)^16,18^, we found that while primed cells certainly showed important differences in their transcriptome profile from non-primed cells, fully resistant cells differed far more dramatically. The expression of key priming marker genes (e.g. *AXL, ITGA3, SERPINE1*) were all considerably higher in resistant cells as compared to primed cells. These results further suggested that primed cells undergo an adaptive process as they become fully therapy-resistant.

To find the regulatory factors that may be involved in this adaptive process, we measured changes in chromatin accessibility as cells became fully therapy-resistant using ATAC-seq^30,31^. We previously showed that chromatin accessibility did not particularly change in primed vs. non-primed cells, but many genomic loci showed changes in chromatin accessibility upon cells becoming fully therapy-resistant^16^. A reanalysis of the motifs in these loci showed enrichment for binding sites for a number of transcription regulators, notably the general transcriptional regulator AP-1 (Fig. 1E). AP-1, which is generally composed of a heterodimer of FOS and JUN subunits, is often identified in ATAC-seq data and disregarded due to its ubiquity. However, our hypothesis is that when a cell transitions from primed to fully resistant, the changes in gene expression are not exclusively predetermined by hard-coded regulatory programming. We reasoned that the very ubiquity of AP-1 would allow it to potentially reprogram cells in a flexible manner depending on the current state of the cell rather than any pre-existing regulatory program.

### Dose escalation demonstrates cellular adaptation during the acquisition of therapy resistance

Much as the expression of most priming marker genes increased during the acquisition of resistance, a hallmark of cellular adaptation, we reasoned that cellular adaptation could also heighten the resistant potential of primed cells. To test this possibility, we used a dose-escalation experimental design that takes advantage of the fact that at lower doses of the therapeutic drug trametinib (1 nM as opposed to 5 nM), about three times as many resistant colonies form. We sought to distinguish selection from cellular adaptation by first treating cells at a low dose for two weeks, then switching to a higher dose for another two weeks, counting the total number of resistant colonies at 4 weeks (Fig. 2A). If resistance were purely a consequence of selection of a preexisting resistant phenotype, then after switching from the lower to the higher dose of therapy, only the original subset of cells that were primed to be resistant to the higher dose to begin with would survive. However, if the cells were adapting during their time in the lower dose of therapy in such a way as to enhance their resistance characteristics, then many more of them would survive upon switching to the higher dose. We found the latter to be the case: with low dose alone for 4 weeks, we measured 67.6 ± 4.9 resistant colonies, high dose alone 20.6 ± 2.0, and 59.3 ± 5.1 resistant colonies when switching from low to high dose, demonstrating that cells are adapting while being exposed to the therapy at low dose (Fig. 2B). It is known that cell density can affect the frequency of therapy resistance, so we also split cells to change their density at the time of dose escalation and found similar results, thus eliminating those effects as potential confounders (Supp. Fig. 1B). We also measured the timescale of the cellular adaptation by exposing cells to the low dose of therapy for different amounts of time before switching to the high dose of therapy, finding that cellular adaptation occurred after around 10-14 days of exposure to the low dose of therapy (Fig. 2C, Supp. Fig. 1C).

**Figure 2:**
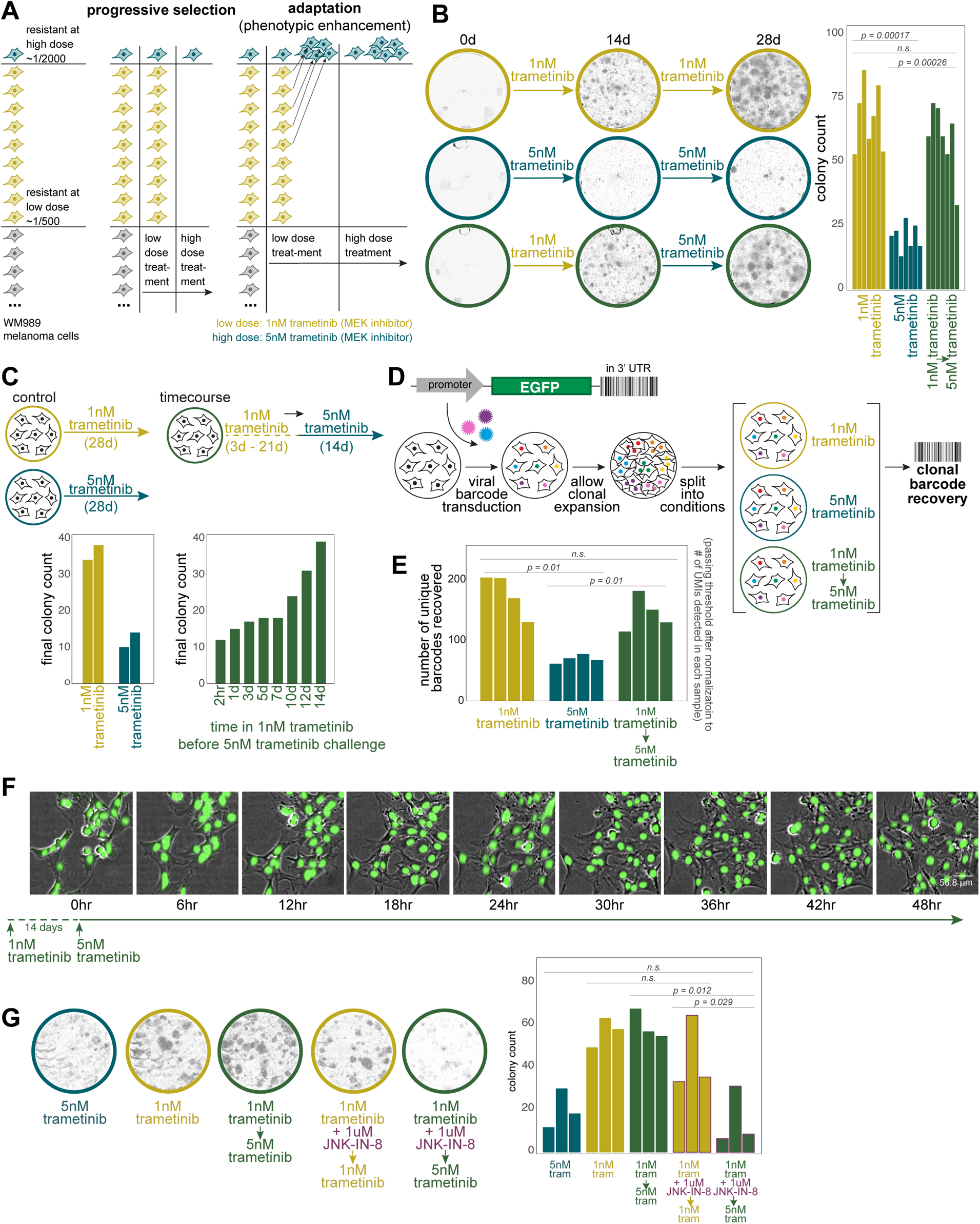
Dose-escalation experiments reveal AP-1-dependent cellular adaptation during the acquisition of therapy resistance. **(A)** Schematic of dose-escalation experimental design to differentiate a purely selective process from an adaptive one. **(B)** Representative images of colonies at approximately day 0 (therapy initiation), day 14 (dose-switch), and day 28 (endpoint) for low dose alone, high dose alone, and low-to-high-dose switch conditions. Quantification of resistant colonies under different treatment conditions. Bar plots show the average number of resistant colonies from 7 biological replicates after 4 weeks of treatment. Error bars represent standard error (SE). Statistical significance was determined using one-way ANOVA followed by Shapiro-Wilk’s test for normality of residuals and Levene’s test for homogeneity of variance. Statistical significance was determined using one-way ANOVA followed by Shapiro-Wilk’s test for normality of residuals and Levene’s test for homogeneity of variance. Post-hoc comparisons were performed using paired t-tests. This statistical approach was applied across all panels in this figure. **(C)** Time course of adaptation. Left: Bar plots comparing the number of resistant colonies under continuous low-dose versus high-dose conditions (two technical replicates in each condition). Right: Bar plots showing the number of resistant colonies when switching to high dose after varying durations of low dose exposure. **(D)** Schematic of DNA barcoding lineage tracking experiment. Cells were barcoded, grown for 4-5 divisions, and then split between 6 arms, each of which was treated as depicted. **(E)** DNA barcode analysis of individual cellular clones. Bar plot showing the number of unique barcodes detected across different treatment conditions. Each bar represents a biological replicate (n=4). Barcode counts are normalized to the number of Unique Molecular Identifiers (UMIs) detected for each sample. **(F)** High-frequency live imaging of a therapy-resistant colony during dose escalation. 48-hour time-lapse images of a colony of H2B-GFP labeled WM989 A6-G3 melanoma cells (WM989 A6-G3-A10) following 14 days of low dose (1nM) trametinib treatment. The first time point shown (0hr) indicates the moment the dose was escalated to 5nM trametinib. Images were captured every 6 hours; images shown are overlays of phase contrast and GFP (nuclei) channels. **(G)** Effect of AP-1 inhibition on cellular adaptation. Left: Representative whole-well images at day 28 (endpoint) showing colony formation under different conditions. Right: Bar plot quantifying final colony counts for each condition. Each bar represents a biological replicate (n=3).

This same effect was also demonstrated by DNA barcoding of individual cellular clones. We barcoded cells and let them divide, then split the twins across the three different conditions (high dose alone, low dose alone, and low-to-high-dose switch). We then harvested all the resistant colonies and sequenced the barcodes to see which ones survived and which twins survived across the various conditions (Fig. 2D) because twins from the same lineage almost always share the same fate^20,32,33^. We found that there were barcodes that specifically survived in the low-to-high-dose switch condition but not in the high dose alone, indicating that these cells were adapting in the presence of the therapy (Fig. 2E). Moreover, our experimental design allowed us to track the fate of twin cells across the various experimental conditions. We found that the twins of the cells that survived in the low-to-high-dose switch condition were also likely to survive in the low-dose alone condition (Supp. Fig. 1D), supporting the idea that it is the additional cells that survive at the low dose that are the ones that survive in the low-to-high-dose switch condition.

Another possibility was that there was progressive selection during the switch from low to high dose—that is, the cells would survive and expand at the low dose of therapy, allowing a small fraction of the colony to then develop resistance to the high dose of therapy. To eliminate this possibility, we monitored cells by high-resolution live imaging during the switch from low to high dose, allowing us to track the fates of individual cells within the colony at the time of dose escalation (Fig. 2F, Supp. Fig. 1E). We found that the vast majority of cells in the colony did not die upon the dose escalation; rather, there was a short pause of 1-2 days in cellular proliferation, after which almost all cells resumed proliferating. Thus, we concluded that the high number of resistant colonies after dose escalation is due to cellular adaptation and not progressive selection of emergent variability.

Previous results from our lab have shown that resistant colonies can come in multiple resistant types^20,34^, raising the possibility that the frequency of these different resistant types was different in the low-to-high-dose switch condition as compared to high or low dose alone. To assess this possibility, we used FateMap (see Goyal et al. 2023 for more details) to quantify transcriptional heterogeneity across resistant colonies in all three conditions (Supp. Fig. 2A). We found that, generally speaking, the distribution of resistant types was similar between the low-to-high-dose switch condition and the low condition, arguing against progressive selection of any particular resistant type (Supp. Fig. 2B). Sometimes, a cell can adopt one type at one therapy dose but adopt a different type at another dose (Supp. Fig. 2C). For 12 barcodes taken as examples, we found that the fate of the barcode from the low-to-high-dose switch condition resembled the fate of its twin in the low condition 8/12 times, high dose 2/12, and all similar 2/12 (Supp. Fig. 2D). These results agree with the notion that cells encode the memory of the initial state within the first two weeks, thus making it more difficult to switch fates once the dose is escalated later.

Given the strong association of AP-1 binding motifs with changes in chromatin accessibility during the transition of primed cells to fully resistant cells, we wondered whether AP-1 was responsible for the adaptive behavior. In the context of the dose-escalation experiments, we hypothesized that AP-1 inhibition would thus lower adaptation potential. We co-treated cells with the Jun N-terminus Kinase (JNK) inhibitor JNK-IN-8 during the low-dose treatment period before switching to the high dose (Fig. 2G). We found that whereas before, many more cells survived the transition to high dose than survived in the high dose alone, with AP-1 inhibition, the rate of survival upon switching from low to high dose was similar to that of continuous high dose alone, showing that cells were no longer able to undergo the adaptive behavior. Importantly, JNK-IN-8 alone did not affect the growth of therapy-naive cells (Supp. Fig. 2E). We saw similar effects using T5224, which inhibits the DNA binding activity of c-Fos/c-JUN (Supp. Fig. 2G). Together, these results show that AP-1 is at least partially responsible for the ability of cells to adapt to therapy.

For a more complete picture of the effects of AP-1 inhibition upon cell fate during the transition to stable resistance, we performed FateMap on resistant colonies as above, but with cotreatment with the AP-1 inhibitor JNK-IN-8. When AP-1 was inhibited, we saw more drift between the cell fates in the low-to-high-dose switch condition as compared to low dose alone, again suggesting that blocking memory formation allowed cells to change their fate rather than locking it in based on the initial state of the cell (Supp. Fig. 2C-D). At the same time, it is important to note that the fact that resistant cells can adopt different fates at low and high doses means that at least some degree of extrinsic conditions affect the ultimate state of the cell^34^, suggesting some degree of a programmed response as well as an encoding based on the initial state of the cell (see Discussion).

### Encoding a memory of an induced response

The above results show that cells change their phenotype during exposure to the therapy. To demonstrate that this phenotypic change was an example of cellular learning, we wanted to show that the specific changes to gene expression during this adaptation were not the result of the execution of a predefined regulatory program but rather consisted at least partly of the encoding of the expression of genes at the time of expression into memory. We reasoned that one way to distinguish between a predetermined cascade of gene expression changes and a response based on memory encoding was to attempt to plant the memory of the expression of a “passenger” gene into primed cells by seeing whether that expression was then committed to memory as the cell became fully resistant.

To demonstrate that the transition from primed to fully resistant cell induces the formation of cellular memories of the state of the cell at the time of therapy initiation, we tested whether we could cause the formation of a memory of the expression of a passenger gene. As an inducer, we used dexamethasone, a steroid that induces expression of a large set of genes through the activation and nuclear translocation of the glucocorticoid receptor^35^, and one that is not generally considered to be a major factor in therapy resistance in melanoma (Supp. Fig. 3A). First, we tested whether treating cells with dexamethasone induced genes in a transient manner, meaning that the removal of the dexamethasone signal would result in a loss of expression of the target gene. Based on bulk RNA sequencing data (Supp. Fig. 3B), we found a set of genes that were induced by dexamethasone and turned off when dexamethasone was removed. Of these, we selected three genes (*FKBP5*, *TGM2*, and *LTBP1*) for further analysis by single-molecule RNA FISH, which enables the counting of transcripts in single cells.

We measured expression induction upon the application of dexamethasone and subsequent reversion after dexamethasone removal. We found that each gene was induced upon 3 days of dexamethasone application, returning to baseline after 14 days of growth post removal. After the reversion time, the cells were quite crowded. To show that the crowding itself was not responsible for the reduced expression after the removal of dexamethasone, we induced cells after crowding to show they were still responsive to dexamethasone (Supp. Fig. 3C). We also split the cells to lower densities after removal and found that expression was still at baseline levels (Supp. Fig. 3E).

Per our hypothesis, if one of these genes were expressed at the point in time that trametinib was applied, its expression would be committed to memory and thus retained even in the absence of the originating dexamethasone signal. We thus applied trametinib after three days of dexamethasone treatment, at which point we also removed dexamethasone. We measured expression in the resultant resistant colonies after 14 days. For *FKBP5*, we found that the expression levels were as high, if not higher, than with dexamethasone alone after 3 days, even though dexamethasone was removed 14 days earlier. Importantly, the addition of trametinib alone did not induce *FKBP5*, showing that the expression of this gene was not inherently associated with therapy resistance (Fig. 3B).

**Figure 3:**
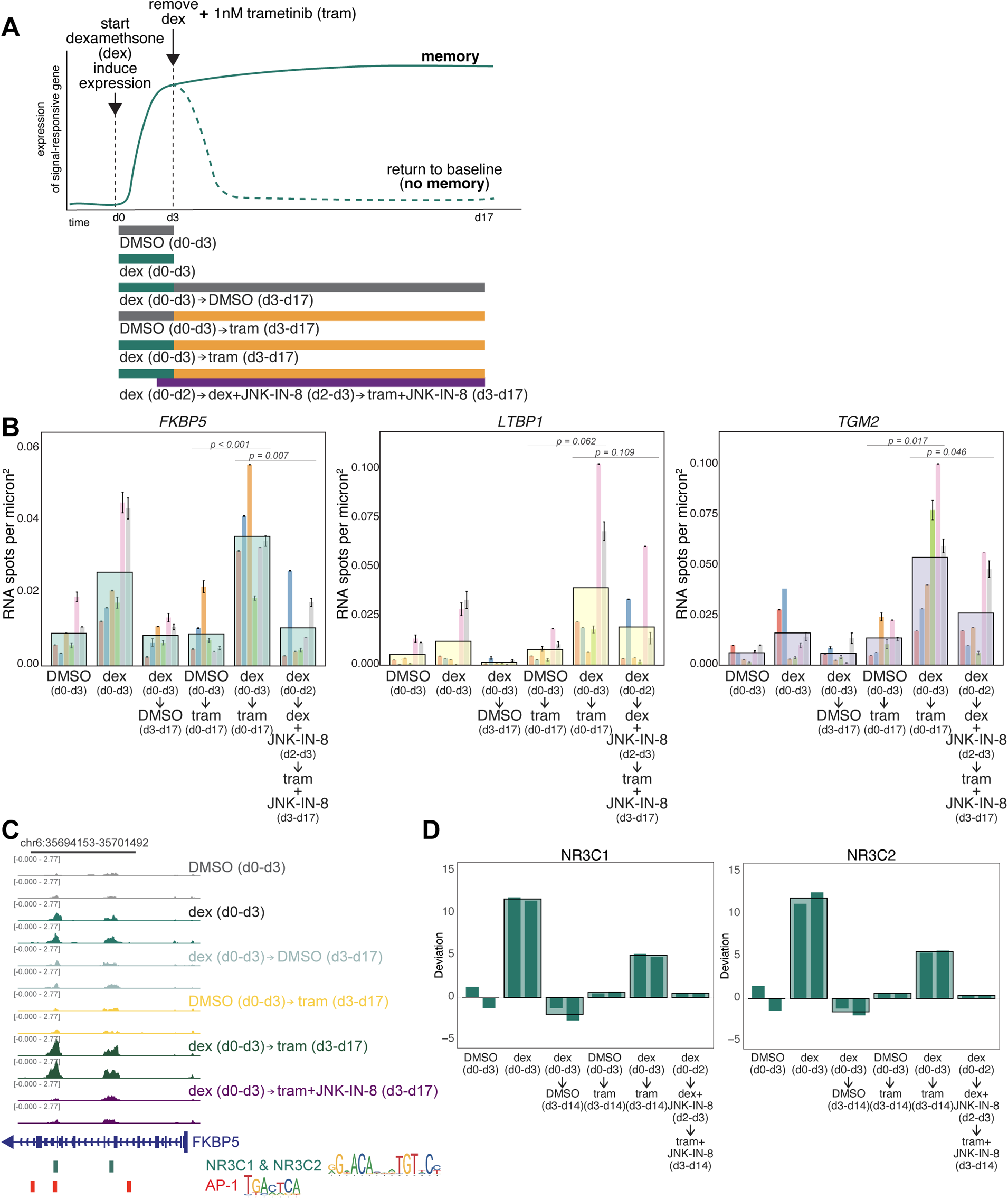
Passenger memory formation in WM989 melanoma cells during resistance acquisition. **(A)** Schematic of the hypothesis that the transition from primed to fully resistant cell induces the formation of cellular memories of the state of the cell at the time of therapy application (top); corresponding experimental design (bottom) **(B)** Single-molecule RNA FISH quantification of *FKBP5*, *TGM2*, and *LTBP1* expression under control and experimental conditions. Large bars represent the mean for each condition, with a total of 602-806 cells analyzed for each condition. Smaller underlying bars show 6 biological replicates, color-coded consistently across conditions and genes. Error bars indicate standard error of the mean (SEM). Statistical significance was determined using one-way ANOVA followed by Shapiro-Wilk’s test for normality of residuals and Levene’s test for homogeneity of variance. Post-hoc comparisons were performed using paired t-tests. **(C)** Example track showing peak changes in conditions described in panel A for the promoter region of *FKBP5*. **(D)** Bar graphs of chromVAR deviation z-scores demonstrating changes in transcription factor signatures for NR3C1(left) and NR3C2 (right). Large bars represent the mean for each condition and smaller underlying bars show the 2 replicates. All experimental conditions shown in Supp. Fig. 4D

For *TGM2* and *LTBP1*, we found a different pattern of expression, albeit one that also indicated memory formation. For these genes, adding trametinib alone did induce expression in the resistant colonies (Fig. 3B). However, treatment with dexamethasone for 3 days beforehand led to stark 4-and 5-fold increases in expression of *TGM2* and *LTBP1*, respectively. This induction level was well beyond that of either trametinib or dexamethasone alone. Thus, for these two genes, the induction of expression was still “memorized” in that without trametinib, the expression level would have returned to baseline, and the expression level was far beyond what trametinib would have induced alone. For all three genes, the expression level at the time of trametinib exposure was committed to memory and affected its future expression, the hallmark of memory and learning. (We noted significant variability between independent biological replicates, likely due to biological variability inherent to cells grown in culture in slightly different conditions. For *TGM2* and *LTBP1*, we found that longer durations of induction resulted in more consistent induction and showed the same memory effect (Supp. Fig. 3E).)

One possible alternative was that once cells become resistant, their induction and reversion timescales upon additional and removal of dexamethasone were different from those of the therapy-naive cells. To test this possibility, we took a population of resistant cells that had never been treated with dexamethasone before and induced them with dexamethasone for 3 days, then cultured them for 7 more days without dexamethasone (Supp. Fig. 3F). We found that the expression of *FKBP5*, *TGM2*, and *LTBP1* all increased upon dexamethasone induction and reverted to baseline levels of expression after dexamethasone was removed. These results show that the mechanism by which these genes are induced is not intrinsically altered by therapy resistance, nor is the sustained expression of these genes merely a consequence of the difference in the regulatory state of resistant cells. Rather, these results show there is a window of opportunity for memory formation during the early period of the addition of trametinib, consistent with our other results.

Our dose escalation results showed that AP-1 is an important factor for adaptive responses, and our prior ATAC-seq data suggested it may play an important role in the regulation of genes that showed enhancement. To test this possibility, we added the JNK inhibitor one day before trametinib treatment (to ensure inhibition of JNK prior to MEK inhibition) and measured the expression levels of *FKBP5*, *TGM2*, and *LTBP1*. We found that the expression of these genes largely reverted back to the levels of trametinib-induction alone. This result shows that the memory encoding can be largely blocked by blocking AP-1 activity. Notably, *FKBP5* expression went back down to baseline (expression in DMSO alone), whereas *TGM2* and especially *LTBP1* went back towards the expression in trametinib alone, suggesting that the expression above and beyond that induced by trametinib was AP-1 dependent, supporting the model that AP-1 is responsible for memory formation.

Building on our observations that AP-1 inhibition blocks memory encoding and reduces enhanced gene expression, we sought to measure changes in chromatin accessibility to identify regulatory factors associated with memory formation. We analyzed changes in chromatin accessibility using ATAC-seq (Fig. 3C-D; Supp. Fig. 4B-D). ATAC-seq yields reads that typically coalesce into peaks that signify open regions of chromatin; these peaks are thought to encompass regions where transcription factors bind. We used chromVAR to compare changes in transcription-factor-associated accessibility as we encoded a memory of an induced response with dexamethasone as in Figure 3A^36^. chromVAR, as applied to our bulk ATAC-seq data here, takes a set of peaks and then tabulates chromatin accessibility data sharing a common feature, such as a specific motif. For each such motif, it calculates a deviation score based on the measured number of associated read fragments compared to a null expectation of chromatin accessibility, normalized by accessibility in a set of background peaks that provide a sense of expected variation. Here, we set the ATAC-seq reads from the 3 days of DMSO conditions as the null expectation from which deviations were calculated.

We first looked at accessibility changes across all peaks detected in samples treated with dexamethasone for three days. As expected, in these samples, peaks containing glucocorticoid receptor (GR; NR3C1 and NR3C2) binding sites exhibited increased chromatin accessibility (Fig. 3C-D, Supp Fig 4B-D). In the conditions where dexamethasone was removed after three days followed by continuous culture in DMSO for an additional 11 days, we saw a decrease in the accessibility of those same sites, showing that chromatin accessibility followed the same transient response as the expression of GR target genes did. Furthermore, trametinib alone did not increase the accessibility of these sites. We then wondered whether the memory encoding effects we saw in the expression of *FKBP5* extended to chromatin accessibility at putative GR binding sites. We saw that in the condition where dexamethasone was removed after three days and trametinib was added for a further 11 days (the memory formation condition), chromatin accessibility stayed high for sites containing GR. These results show that trametinib treatment reinforced the chromatin accessibility changes that were initially induced by dexamethasone, potentially facilitating the observed memory effects on expression.

**Figure 4:**
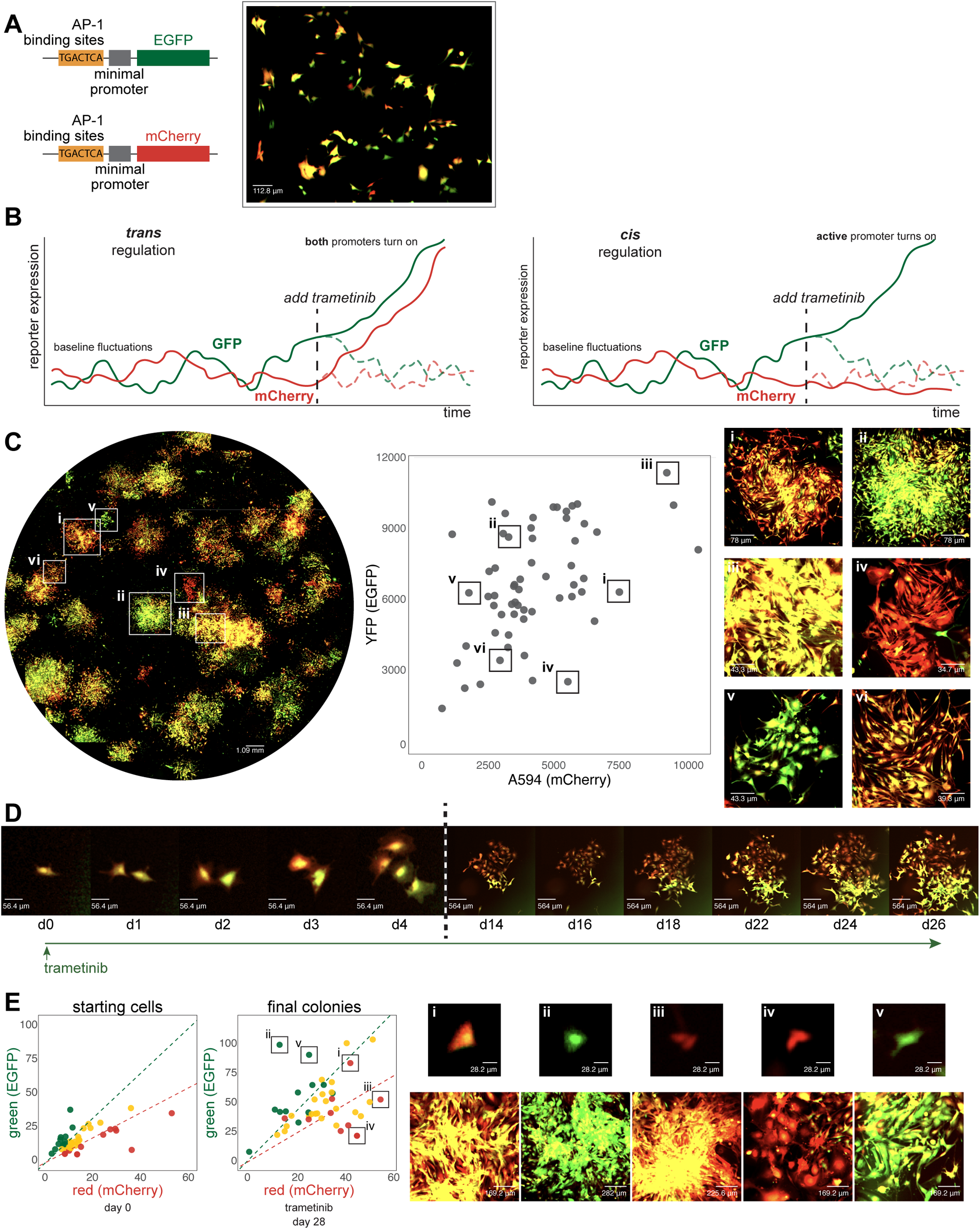
Characterization of AP-1 activity dynamics and memory in WM989 melanoma cells using a dual-color reporter system. **(A)** Left: Diagram of two lentiviral constructs containing AP-1 binding sites (6 palindromic repeats of TGACTCA) upstream of a minimal promoter driving expression of either EGFP (top) or mCherry (bottom). Right: Image showing cell-to-cell variability in expression in a single-cell-derived clone of WM989 A6-G3 cells transduced with both reporter constructs (WM989 A6-G3-BE5) In addition to the BE5 line, we also isolated and validated 3 additional lines; CC10, DE5, AB1. **(B)** Diagram comparing potential observations in this reporter system under two scenarios: sequence-dependent regulation (left) versus activity-dependent regulation (right) of memory. **(C)** Left: Stitched 10x magnification image of a well containing WM989 A6-G3-BE5 AP-1 reporter cells treated with 1nM trametinib for 28 days. Middle: Scatter plot of fluorescence intensities of colonies formed in the image to the left (calculated as the 90th percentile intensity of each colony minus the mean of the median intensities of randomly selected areas without colonies). Right: Zoomed-in images of 6 selected colonies (i-vi) demonstrated variation in mCherry and EGFP ratios and intensities. These 6 colonies have been identified and labeled on both the scatter plot and the stitched well image. **(D)** Time-lapse imaging of the formation of a resistant colony from WM989 A6-G3-BE5 AP-1 reporter cells over 26 days under treatment with 1nM trametinib (starting at day 0). Images from the first 4 days (d0-d4; left of the dotted line) have the same magnification, and images from the final 12 days (d14-d26; right of the dotted line) all have 10x lower magnification as the cells’ images on the left. Images were contrasted so that the background appears black for ease of visual comparison. **(E)** Left: Scatter plots of fluorescence quantifications from Incucyte S3 images of WM989-A6-G3-BE5 cells treated with 1 nM trametinib for 28 days. Colonies were traced back through time-lapse images to identify their originating cells. The initial red-to-green fluorescence intensity ratio of these starting cells was used to categorize the initial ‘color’ of the cell (quantified as the mean of the cell minus the median of the 10 px annulus around the cell): top 25th percentile as red, bottom 25th percentile as green, and the remainder as yellow. This initial categorization is used to determine the color coding of the final colonies. Right: Zoomed-in images of 5 selected colonies (i-v) have been identified and labeled on the final colony scatter plot (in order to account of spaces between cells within a colony, colony fluorescence is quantified as the mean fluorescence of the pixels in the annotation above the median of the colony annotation, minus the median of the colony annotation). Both the starting cell (top) and the resistant colony arising from that cell (bottom) are shown.

Given our results showing that inhibition of AP-1 can block memory formation, we asked whether AP-1 inhibition also prevented these reinforcements in chromatin accessibility due to trametinib exposure. As in the memory encoding experiment in Figure 3A-B, we treated cells with dexamethasone for three days, then exposed them to trametinib while removing dexamethasone, adding the JNK inhibitor on the final day of dexamethasone treatment and continuing it throughout the trametinib treatment period. We found that JNK inhibition blocked the increase in the chromatin accessibility of GR-sequence-containing regions that we had seen in the presence of trametinib without JNK inhibition; indeed, the levels of chromatin accessibility went down almost to baseline levels, suggesting that AP-1 activity is critical for maintaining GR-associated chromatin accessibility. Of note, regions containing AP-1 binding motifs did not show significant accessibility changes in any condition where trametinib was added. Given our previous results^16^, we suspect that the signature would emerge after a longer time in trametinib than the 11 days we used here.

### AP-1 encodes memories in *cis*

As outlined earlier, epigenetic memory can be stored either in *cis* or *trans*^8^, with the former referring to regulatory modifications at the genomic locus (such as DNA methylation or histone modification) and the latter referring to the state of the cell composed of the levels and activity of transcription factors and other regulatory molecules. An important practical distinction is that *trans* epigenetic memory is encoded in the hard-wired interactions between various molecules (such as transcriptional feedback loops), which are ultimately determined by the genetic sequence itself, whereas *cis* memory is encoded more flexibly in ways that are not tied to the sequence intrinsically.

We thus wondered whether the epigenetic memory in our system was encoded in *cis* or *trans*. To do so, we first constructed a genetic reporter that would enable defined, synthetic epigenetic memory formation. Given that AP-1 was important for the formation of these memories, we designed a reporter system in which six palindromic repeats of the canonical AP-1 binding site TGACTCA were placed immediately upstream of a minimal promoter that drove the expression of either EGFP or mCherry. We transduced each reporter into a population of WM989 A6-G3 melanoma cells and isolated four clonal cell lines in which both the EGFP and mCherry reporters were integrated. We verified the responsiveness and specificity of the reporters by showing that expression was induced with PMA (an inducer of AP-1) and was reduced when treated with JNK-IN-8 (Supp. Fig. 5A). Also, reporters without AP-1 binding sites were not responsive to induction (Supp. Fig. 5D).

In therapy-naive cells at baseline, fluctuations in gene expression led to variability in the expression of the two reporter genes in individual cells^37,38^, leading some cells to have more EGFP expression, some to have more mCherry expression, and some to have both (Fig. 4A). The autocorrelation time-scales of the fluctuations were 17.4 hours and 18.8 hours for EGFP and mCherry, respectively, and the cross-correlation time-scale was 8 hours, showing that there was relatively limited memory in the system before the application of therapy (Supp. Fig. 5B-C).

Based on our previous results, we anticipated that the expression of the reporter gene would be committed to memory over time after the initiation of therapy. The dual reporter setup allowed us to distinguish whether this expression commitment would be due to regulation in *trans* (meaning regulation by sequence-specific binding of AP-1 to its binding sites) or in *cis* (in our case, meaning regulation by association; i.e., dependent on the existing expression levels at the time of therapy initiation). In the *trans* case, both reporters would be committed to memory by AP-1 activity because they have identical regulatory sequences. However, in the *cis* case, given the baseline fluctuations, we would expect that if a primed cell had higher instantaneous expression of the EGFP reporter at the time of therapy initiation, then the resultant resistant colony would have higher levels of EGFP than mCherry and vice versa (Fig. 4B). We formed resistant colonies in this dual reporter cell line and observed the latter: resistant colonies showed a range of levels of EGFP and mCherry, with some appearing almost exclusively EGFP-positive, some mCherry-positive, and some both (Fig. 4C). To confirm that the reporter expression was AP-1 dependent, we added JNK-IN-8 concurrently with resistant colony formation and found that the expression of the reporter was greatly attenuated (Supp. Fig. 6B-C).

To demonstrate explicitly that the initial state of the cell was encoded into the epigenetic memory, we performed time-lapse tracking of cells through the process of colony formation. We quantified the fluorescence levels of the initial state of the cell and then connected those cells to the ultimate therapy-resistant colony. An example is shown in Figure 4D, in which a single cell gives rise to two cells, one of which expressed more mCherry while the other expressed more EGFP. The resulting therapy-resistant colonies reflected these initial imbalances. Looking across multiple such instances (46 tracks in total), we found that cells that were initially more mCherry or EGFP-positive ended up forming resistant colonies that largely preserved those differences (Fig. 4E). JNK inhibition led to a reduction in the induction of the levels of both reporters and also led to a loss of preservation of their relative levels (Supp. Fig. 6B-C). These results show that probabilistically-arising expression of genes can be encoded into *cis* epigenetic memory in an AP-1-dependent manner.

We also wondered how durable the memory was over time and whether its retention was dependent on AP-1. We isolated five individual therapy-resistant colonies that harbored both copies of the reporter and grew them in trametinib continuously for a month. They continued to maintain high levels of expression of both fluorescent proteins. We then grew three of these individual therapy-resistant colonies (A2, BA1, C3) with and without trametinib and with and without JNK inhibitor for another 7 days. We found that the signal was still largely present in all cases, with a few cells losing expression in the presence of the JNK inhibitor (we also noticed a growth rate defect in the JNK inhibitor alone condition). These results suggest that once the epigenetic memory is formed, it is no longer dependent on either the effects of trametinib or the activity of AP-1, consistent with the possibility that another molecular mechanism is in place in *cis* to maintain the memory (Supp. Fig. 6D).

Together, these results demonstrate that the epigenetic memory formed by the addition of trametinib is encoded in *cis*, supporting the regulation by association model instead of regulation by sequence, and that AP-1 is both necessary and sufficient for its formation. Additionally, these results show that the system is capable of encoding memories of probabilistically arising signals and not just the induced signals in Fig. 3.

## Discussion

Here, we have shown that in the context of the acquisition of therapy resistance, cells are able to learn, meaning that they are able to form memories of their current state at the time of exposure to therapy, and those memories can influence their future behavior. We demonstrated the formation of memories of both induced and probabilistically arising expression of genes. AP-1 emerged as a critical regulator of this process.

Our work suggests that cells have the capability of altering their regulatory networks in ways that are not strictly specified by their intrinsic coding. The generally accepted framework is that regulatory networks are shaped by precise, hard-coded molecular interactions—for instance, the interaction between a transcription factor and its binding site. We show that cells are able to express genes that are not explicitly a part of any existing regulatory network, and the process of committing those genes’ expression to memory is not based solely on the regulatory sequence but also on the state of the genes’ activation^4^. This “regulation by association” is a potentially powerful way for cells to adapt to new situations.

It is possible for conventional regulation and cellular learning to occur simultaneously. Indeed, there is evidence for both in the therapy-resistance model system described here, which comes from tracing the fates of individual primed cells into the variety of resistant types we have previously detected^20^. Those results showed that “twin” cells subjected to the same dose of therapy adopted remarkably similar fates, showing that resistant type specification was strongly intrinsically determined—indeed, twin resistant colonies were far more similar than any other two non-twin resistant colonies, consistent with memory encoding of an initial intrinsic state that is virtually the same in twin primed cells but different in non-twin primed cells. At the same time, we also observed that when twin cells were exposed to two different doses, sometimes the ultimate resistant type would be different between these doses, suggesting that the nature of the challenge posed to the primed cells is an extrinsic influence on its fate. The relative contribution of intrinsic memory formation and extrinsic influence on the trajectory of the cell likely varies by context.

Our work shows that these memories are formed in *cis*, meaning that the molecular encoding of the memory happens at the physical location of the gene itself. This *cis* encoding must be governed by activating molecular epigenetic signals. Most *cis* epigenetic signals are negative regulators, meaning that they serve to form memories of repression, for example, through DNA methylation or the initiation and propagation of repressive histone modifications like H3K9me3 and H3K27me3^39,40^. In those cases, enzymes that read and write such modifications are the key to their epigenetic propagation, such as SUV39H1/2^40^ and PRC2^41–43^, respectively. Importantly, the reading of the epigenetic mark promotes the writing of the same mark, thus propagating the memory. There are, conversely, few, if any, examples of activating epigenetically propagated marks^44^. However, many molecular complexes have bromodomains that bind to activating marks via bromodomains while also being able to deposit such marks, such as CBP/p300, which both binds to acetylated H3K27^45,46^ and acetylates H3K27. Another possible mechanism of memory encoding is chromatin remodeling, in which histone-DNA interactions are modified or nucleosomes repositioned by BAF/BRG1; AP-1 is known to interact with such factors as well^47,48^. Thus, these complexes have the potential to be the basis of activating epigenetic memory. Regulation by nuclear pore proteins have also been shown to propagate memory of prior gene activation, leading to faster reactivation^12,15^. Further work in this area will be required to determine what modifications, enzymes, and feedback interactions are at play in the formation of the epigenetic mechanisms we have outlined here.

We have focused here on epigenetic memory formation in primed cells, i.e., those that become therapy resistant. It remains unclear whether every cell in the initial therapy-naive population can form such memories or if only the rare cells primed for therapy resistance are. It is also possible that there is some intermediate subpopulation that is larger than the primed subset that is capable of memory formation. In our system, it is difficult to distinguish these possibilities because the memory-inducing agent (in this case, therapy) is also a selective agent that kills the non-primed cells that comprise the vast majority of the population. Non-selective means of triggering memories may specifically reveal the memory-forming subpopulations. The characterization of this subpopulation will help show what specific features are required in order for a cell to encode a memory.

AP-1 is downstream of many stress response pathways^49^, and so one possibility is that the stress induced by the response to trametinib is responsible for the encoding of memory. Similar memory-inducing effects tied to AP-1 have been observed in wound healing, inflammatory responses, and aging^7,50–52^, and it may very well be that the same mechanisms are at play. Several studies have also shown that stress, in general, can lead to epigenetic changes, often via heat-shock pathways^53,54^. *Cis*-encoded epigenetic memories may allow cells to respond to complex stimuli and situations that they do not have an existing regulatory program to handle.

One consequence of stress-induced memories is the potential for the accidental formation of deleterious memories. A cell, upon encountering stress, may form a memory of a transiently expressing a gene whose expression is in some way incompatible with or otherwise harmful to the cell. The accumulation of deleterious memories may lead cells to function less well over time, constituting a mechanism for the generation of epigenetic damage during aging. Removing such memories from cells could potentially restore their prior functionality. In the context of therapy resistance in cancer, removing such memories may cause resistant cells to revert to a cellular state that is now vulnerable to either the original therapeutic agent or some new set of agents.

We have here drawn a distinction between programmed and learned responses to stimuli (*cis* vs. *trans* epigenetics, respectively). In the most general terms, the distinction is artificial in the sense that even responses based on learning can be thought of as the execution of a program. We think a more philosophically coherent description is to consider the mapping between the inputs to and outputs from the cell. Program execution corresponds to what could be considered an input funnel, in which many different inputs lead to the same output (say, a developmental outcome as a result of a regulatory cascade). Learning is more akin to a one-to-one mapping, in which each different input leads to a different output. Such output contingency is a hallmark of learning and is a practically useful distinction.

## Acknowledgments

We thank the Raj lab members for scientific discussion and comments on the manuscript, particularly Karun Kiani, Sam Reffsin, Grant Kinsler, Catherine Triandafillou, Yael Heyman, Laura Van Eyndhoven, Miles Arnett, and Vinay Ayyappan; the Genomics Facility at the Wistar Institute, especially S. Majumdar and S. Widura, for assistance with sequencing and single-cell partitioning and addition of 10X cell identifiers; the Flow Cytometry Core Laboratory at the Children’s Hospital of Philadelphia Research Institute for assistance with flow cytometry and fluorescence-activated cell sorting; and the William Greenleaf lab for sharing their expertise in ATAC-seq and providing us with the custom primers used in our ATAC-seq experiments; professors Marissa Bartolomei, Roberto Bonasio, John Murray, and Rajan Jain for generously sharing their time and expertise to comment on the manuscript.. A.R. acknowledges support from a center grant from the Mark Foundation for Cancer Research, NIH Director’s Transformative Research Award R01 GM137425, NIH R01 CA238237, NIH R01 CA232256, and NIH 4DN U01 DK127405. J.L., P.T.R., R.H.B., and N.J. acknowledge support from NIH Medical Scientist Training Program T32GM007170. R.H.B. acknowledges support from NIH Training Grant In Computational Genomics T32HG000046. G.T.B. and A.O. acknowledge support from NSF GRFP DGE-2236662. Z.N. acknowledges support from the Roy and Diana Vagelos Scholars Program in the Molecular Life Sciences, Roy and Diana Vagelos Science Challenge Award, Barry M. Goldwater Scholarship, Fannie and John Hertz Foundation Fellowship, Paul & Daisy Soros Fellowship for New Americans, and Department of Energy Computational Science Graduate Fellowship. K.S. acknowledges support from NIH R01 GM143229. Y.G. acknowledges support from the Burroughs Wellcome Fund Career Awards at the Scientific Interface and Cancer Research Foundation Young Investigator Award.

## Author contributions

A.R. conceived the project. A.R. and J.L. designed all experiments. P.T.R. assisted J.L. with bulk RNA-seq experiments and analysis. A.O. assisted J.L. with ATAC-seq experiments and P.T.R. performed analysis with input from J.L. and A.R. K.S. provide the Tn5 used in all ATAC-seq experiments. Z.N. developed the algorithm for automated spot detection used for sm RNA FISH analysis and S.W. and M.G.D. assisted with the manual segmentation used in the analysis of sm RNA FISH data. N.J. and Y.G. assisted with barcode library preparation for both the DNA lineage tracing and FateMap experiments, and N.J. provided the computational pipeline used in the analyses. Y.G. assisted J.L. with DNA lineage tracing and FateMap experiments. G.T.B. assisted J.L. with the manual isolation of resistant cell lines. J.L., P.T.R, and A.R. prepared all illustrations used in this study. A.R. wrote the manuscript with assistance from J.L. and P.T.R. and with input from all authors.

## Declaration of generative AI and AI-assisted technologies

During the preparation of this work, the authors used chatGPT and Claude to generate/comment code for analyses. After using this tool or service, the authors reviewed and edited the content and take full responsibility for the content of the publication.

## Supplemental information

Document S1. Figures S1–S6 and Tables 1-4.

## Methods

### Cell culture

WM989 melanoma cells from the laboratory of Meenhard Herlyn were validated by DNA short tandem repeat (STR) microsatellite fingerprinting at the Wistar Institute. These cells were cultured in TU2% media containing 78% MCDB (Sigma M7403), 20% Leibovitz’s L-15 media (Life Technologies Inc., 11415064), 2% FBS, 1.68 mM CaCl2, and 50 U ml-1 penicillin and 50 μg ml-1 streptomycin (Invitrogen 15140122). WM989-A6-G3 cells, first described in^16^, are specific single-cell-derived subclones of WM989^28^ that were used to minimize genetic heterogeneity. WM989 A6-G3 AP1-dualColorReporter cells, including lineages DE5 and CC10 are further single-cell subclones of the original WM989 A6-G3 lineage and cultured in the same conditions. WM989 cell lines were passaged using 0.05% trypsin-EDTA (Invitrogen 25300054). All cell lines tested negative for mycoplasma. Cells were cultured in Corning Falcon plates (12-well: 08-722-29, 24-well: 08-722-1) for all Incucyte S3 time-lapse experiments and on Cellvis 24-well glass bottom plates (Fisher Scientific NC0397150) for RNA FISH.

### Drug Treatment

We prepared stock solutions in DMSO of 50 μM trametinib (Selleckchem S2673), 10 mM vemurafenib (PLX4032) (Selleckchem S1267), 250 μM PMA (Phorbol 12-myristate 13-acetate) (Selleckchem S7791), 10 mM JNK-IN-8 (Selleckchem S4901), and 10 μM T-5224 (Selleckchem S8966). We prepared small aliquots (10-20 μl) for each drug and stored them at −80 °C to minimize freeze-thaw cycles. For drug treatment experiments, we diluted the stock solutions in culture medium to final concentrations of 5 nM and 1 nM for trametinib, 100 nM and 1 μM for vemurafenib, 250 nM for PMA, 1 μM for JNK-IN-8, and 1 μM for T-5224, unless otherwise specified. The doses of trametinib and vemurafenib were chosen based on a dose curve to obtain virtually complete growth arrest. Doses of JNK-IN-8 and T-5224 were chosen based on dose-testing for which treatment with JNK-IN-8 or T-5224 alone had no significant impact on growth rate. Medium was replaced every 3-4 days for all experiments. At the end of the treatment, surviving cells were trypsinized, neutralized, washed with 1× DPBS, and then either (1) pelleted and stored at −20 °C for gDNA extraction, (2) resuspended in PBS for scRNA-seq experiments and ATAC-seq experiment, (3) lysed with QIAzol and stored at -80 °C for bulk RNA sequencing, or (4) fixed for imaging at the end of the treatment as per the protocol detailed below.

### Manual isolation of resistant colonies

For manual isolation of resistant colonies, cells were plated at a density of 20,000 cells on 15-cm tissue culture plates to prevent overlap of colonies and grown in 1nM trametinib for 21 days. After this period, plates with resistant colonies were scanned under a tissue culture microscope to identify physically isolated colonies.

These distant colonies were physically isolated and dissociated via treatment with 0.05% trypsin for 5–10 minutes, with dissociation time varying between colonies. Colony suspensions were then plated in 96-well plates containing 180 μl of TU2% medium with 1 nM trametinib. The following day, 150 μl of medium was replaced to remove residual trypsin without disturbing adherent cells. Isolated resistant colonies were closely monitored daily for growth, and the 1nM trametinib-containing medium was changed every 3–5 days. As the isolated resistant colonies reached 70–80% confluence, they were serially expanded into 12-well, 6-well, and then 10-cm plates. This procedure ensured the isolation and expansion of individual resistant colonies for further analysis.

### Time-lapse microscopy

Time-lapse experiments were conducted on an Incucyte S3 Live Cell Imaging Analysis System (Sartorius) with a 4× objective. For phenomenology experiments, only phase imaging was collected at 4x magnification. For fluorescent imaging of AP-1 reporter cells, with 300ms exposure for GFP and 400ms exposure for mCherry approximately every 24 hours unless otherwise specified. For higher-resolution time-lapse imaging of resistant colonies during high-dose trametinib (5 nM) challenge, we used WM989 A6-G3 cells tagged with an EGFP nuclear reporter (H2B-EGFP). Images were captured every 6 hours to enable precise tracking of individual cells within colonies. Images were exported as 8-bit TIFs for phase images and 16-bit TIFs for fluorescent images. These were processed using a custom script (https://github.com/arjunrajlaboratory/process_incucyte_tiff_data) prior to upload and analysis with NimbusImage (see Image analysis section for more details).

### Flow sorting

We used 0.05% trypsin-EDTA to detach barcoded cells from the plate and subsequently neutralize the trypsin with the corresponding medium. We then pelleted the cells, performed a wash with 1× DPBS, and resuspended them again in 1× DPBS. Cells were sorted on a BD FACSJazz machine (BD Biosciences) or MoFlo Astrios (Beckman Coulter) with a 100 μm nozzle, gated for positive GFP signal and singlets. Sorted cells were then centrifuged to remove the supernatant medium containing PBS, and replated with the appropriate cell culture medium.

### Fluorescent reporter generation

For WM989 A6-G3 AP1-dualColorReporter cell lines, we adapted the original retroviral construct ^55^ by cloning the mCherry reporter construct with minimal promoter preceded by 6 palindromic repeats of AP1 upstream of the mCherry fluorophore into a lentiviral backbone (pLV-EGFP) by PCR amplification and gel purification (NEB T1020S) of the sequence of interest. PCR primers contained overhangs of a ClaI site at the 5’ end and a SalI site at the 3’ end. Both the lentiviral plasmid and the PCR amplicon were double digested with ClaI (R0197S) and SalI (R3138S) and processed with a PCR & DNA cleanup kit (NEB T1030S). To generate the EGFP reporter, we PCR amplified the AP1 and promoter region of the original mCherry reporter with overhangs to provide cut sites for ClaI and AgeI on the 5’ and 3’ ends, respectively. We then performed a double digest of both the PCR amplicon and the original lentiviral plasmid with ClaI and AgeI. These linearized plasmids with sticky ends were ligated to their corresponding ap1 promoter and lentiviral backbone pairs with T4 DNA ligase (NEB M0202S) and transformed into STBL3 competent cells (Invitrogen C737303) for amplification and plasmid DNA isolation (Qiagen 12362). To generate the clonal lines of the reporter cells, we first transduced WM989-A6-G3 cells, as detailed below. We then isolated fluorescent cells by FACS and generated clonal cell lines by serial dilution. To confirm that both mCherry and EGFP fluorescence were responsive to changes in AP1 activity, we compared DMSO, JNK-IN-8, and PMA. To verify that these changes in fluorescence were specifically driven by the AP1 binding sites, we also generated control plasmids without the AP1 binding sites by the same methods described above, except the 5’ PCR primer was designed to skip the region containing AP1 repeats. These cells were also sorted and cloned as the reporter cells and subject to the same test conditions. We confirmed that the AP1 reporter responded as predicted, while the control did not have any notable response (Supplemental Figure 5D). PCR primer sequences for all reactions above are documented in Supplemental table 1. All plasmid sequences were confirmed through full plasmid sequencing by Plasmidsaurus and are available here: https://benchling.com/jess_li_ljx/f_/RX0xoDXY-ap-1-reporter-plasmids/.

### Single-molecule RNA FISH

We designed custom oligonucleotide probe sets complementary to our genes of interest using custom MATLAB software (available at https://flintbox.com/public/project/50547/) and ordered them from IDT (probe sequences available in Supplemental Table 3). We pooled 32 oligonucleotides targeting each gene and an amine group was added on the 3’ end with TdT. For the 3’ TDT addition, we first separately resuspend Amino-11-ddUTP (Lumiprobe A5040) to 10 mM and IDT oligos to 100 μM in nuclease-free water (NF H_2_O). We then aliquoted 5 μL of each oligo in the probe set into a 1.5mL Eppendorf tube for each corresponding gene. For each probe set containing 32 oligos, we added 4 μL of the Amino-11-ddUTP (for a molar excess of 2.5), 24 μL of 10x TDT buffer, 47.2 μL of 2.5 mM CoCl_2_, and lastly 4.8 μL of TDT (NEB M0315L). We incubated the reaction in a thermocycler at 37°C (lid at 50°C) overnight before proceeding to precipitation. We precipitated the probes with 120 μL of 7.5 M ammonium acetate and 1.08 mL of 190 proof ethanol at -80°C for 30 minutes and spun down the tube at 4°C for 10 minutes at 16,000 x g. The supernatant was removed and the pellet was washed with cold (−20°C) freshly prepared 80% ethanol before spinning down again at 4°C for 5 minutes. We again removed the supernatant and then spun for an additional minute. After this final spin, the tube was opened and left to dry at room temperature for 30 minutes before coupling each set to either Cy3, Alexa 594, Atto647N, or Atto 700. Single-molecule RNA FISH was performed as described in ^56^. In brief, cells were fixed in 4% formaldehyde for 10 minutes at room temperature, permeabilized in 70% ethanol, and stored at 4°C. For hybridization, samples were washed with buffer containing 10% formamide and 2x SSC, then incubated with hybridization buffer containing custom RNA FISH probes, 10% formamide, 2x SSC, and 10% dextran sulfate. Samples were hybridized overnight at 37°C, followed by two 30-minute washes at 37°C. For imaging, cells were DAPI stained and transferred to 2x SSC. Samples were imaged on a Nikon Ti-E with a 60X Plan-Apo objective and filter sets for DAPI, Cy3, Atto647N, Alexa594, and Atto700. Nikon Elements imaging software was used to acquire tiled image grids. The Nikon Perfect Focus System ensured images remained in focus. All images were acquired at 60x magnification and 1.4 numerical aperture.

### Image analysis

Image analysis was conducted using NimbusImage (https://github.com/Kitware/UPennContrast) unless otherwise noted. Cell segmentation was performed manually due to irregular cell shapes and overlapping in resistant colonies. For colony segmentation, we used SegmentAnything (https://github.com/facebookresearch/segment-anything), a built-in function of NimbusImage. Colonies were tracked across time points throughout the experiment to distinguish between single-cell-derived colonies and merged colonies. For time-lapse analysis of naive cells, SegmentAnything was used to annotate individual cells, tracking them every 8 hours. Fluorescence intensity calculations varied based on imaging conditions. For 10x magnification images (Nikon scope), colony fluorescence was calculated as the 90th percentile of the colony’s fluorescence minus the median fluorescence of randomly selected colony-free areas. For 4x magnification images (Incucyte S3), individual cell fluorescence was calculated as the mean fluorescence of the cell annotation minus the median fluorescence of a 10-pixel annulus around the annotation, while whole colony fluorescence was calculated as the mean of all pixels above the median fluorescence within the colony annotation, minus the annotation’s median fluorescence. Bulk fluorescence intensity analysis for AP-1 reporter controls imaged on Incucyte S3 was conducted directly on the Incucyte S3 software by calculating the fluorescence intensity (GCU or RCU per well, for green and red signals, respectively), normalized to the % confluency of the well. For sm RNA FISH experiments, RNA spots were identified using Piscis, a deep-learning algorithm (https://github.com/zjniu/Piscis)^57^. Spot counts were normalized to the cell area and displayed as spots per micron². In densely packed cell cases, Cellpose2 was used to segment DAPI-stained nuclei and Piscis-detected spots were associated with the nearest nucleus, displayed as spots per nucleus. Image annotation data were exported as CSVs for further processing and graphing in R. NimbusImage annotations and connections were also exported in JSON format.

### Lineage Barcode library construction

Barcode libraries were constructed as previously described, and the protocol is available at https://www.protocols.io/view/barcode-plasmid-library-cloning-4hggt3w. In brief, we modified the LRG2.1T plasmid (gift from J. Shi) by removing the U6 promoter and single guide RNA scaffold and inserted a spacer sequence flanked by EcoRV restriction sites after the stop codon of GFP, subsequently digesting this vector backbone with EcoRV (NEB) and gel purifying the linearized vector. We ordered PAGE-purified ultramer oligonucleotides (IDT) containing 100 nucleotides with a repeating WSN pattern (W = A or T, S = G or C, N = any) surrounded by 30 nucleotides homologous to the vector insertion site. We subsequently used Gibson assembly followed by column purification to combine the linearized vector and barcode oligonucleotide insert. We performed nine electroporations of the column-purified plasmid into Endura electrocompetent Escherichia coli cells (Lucigen) using a Gene Pulser Xcell (Bio-Rad). We then allowed for their recovery before plating serial dilutions and seeding cultures for maxi-preparation. We incubated these cultures on a shaker at 225 rpm and 32 °C for 12–14 h, pelleted the resulting cultures by centrifugation, and used the EndoFree Plasmid Maxi Kit (Qiagen 12362) to isolate plasmid according to the manufacturer’s protocol. Barcode insertion was verified by polymerase chain reaction (PCR) on colonies from plated serial dilutions. We pooled the plasmids from the 9 separate cultures in equal amounts by weight before packaging into lentivirus.

### Lentiviral packaging and transduction

We adapted previously described protocols to package lentivirus^58^. We first grew Lenti-X-293T cells (Takara 632180) to near confluency (75-90%) in 10-cm plates in DMEM containing 10% FBS and 50 U ml−1 penicillin, and 50 μg ml−1 streptomycin, and one day before plasmid transfection, we changed the medium in Lenti-X-293T cells to DMEM containing 10% FBS without antibiotics. For each 10-cm plate, we added 80 μl of polyethylenimine (Polysciences 23966) to 500 μl of Opti-MEM (Thermo Fisher 31985062), separately combining 5 μg of VSVG and 7.5 μg of pPAX2 and 7.35 μg of the barcode plasmid library in 500 μl of Opti-MEM. We then incubated both solutions separately at room temperature for 5 min. We then mixed both solutions together by vortexing and then incubated the combined plasmid–polyethylenimine solution at room temperature for 15 minutes. We added 1.09 ml of the combined plasmid–polyethylenimine solution dropwise to each 10-cm dish. After 6–7 h, we aspirated the medium from the cells, washed the cells with 1× DPBS, and added fresh TU2% medium. The next morning, we aspirated the medium and added fresh TU2% medium.

Approximately 9–11 h later, we transferred the virus-laden medium to an empty, sterile 50-ml tube and stored it at 4 °C, and added fresh TU2% medium to each plate. We continued to collect the virus-laden medium every 9–11 h for the next ∼30 h in the same 50-ml tube and stored the collected medium at 4°C. Upon final collection, we filtered the virus-laden medium through a 0.45-μm PES filter (MilliporeSigma SE1M003M00) and stored 1.5-ml aliquots in cryovials at −80 °C. To transduce WM989 A6-G3 cells, we freshly thawed virus-laden medium on ice, added it to dissociated cells, and plated ∼100,000 cells per well in a 6-well plate with ∼3 ml of the medium. We then incubated the 6-well plate at 37 °C and replaced the medium at ∼8 h, washed with 1× DPBS, and added fresh medium After ∼24 h, we passaged the cells to 10-cm dishes, at which point we typically combined 2 wells by plating them together in a 10-cm dish. For the FateMap experiments with WM989 A6-G3 melanoma cells, we planned to start each split with 500,000 barcoded (GFP-positive) cells. The barcoded cells (GFP-positive) were then sorted and plated for a total of 4–5 population doublings before trypsinization and plating in separate experimental conditions. The volume of the virus-laden medium was decided by performing titration on each cell line and target MOI. We targeted an MOI of ∼10–25% to minimize the fraction of cells with multiple unique barcodes.

### Lineage barcode library sequencing

We followed the protocol originally described in^18^ for lineage barcode library construction and sequencing Genomic DNA (gDNA) was isolated from barcoded cells using the QIAmp DNA Mini Kit (Qiagen 51304) according to the manufacturer’s protocol. We performed targeted amplification of the integrated barcode vector using custom primers containing Illumina adapter sequences, unique sample indexes, variable length staggered bases, and 6 random nucleotides ("UMI"; NHNNNN). For each sample, we performed multiple PCR reactions (using 20–40% of the total isolated gDNA), each consisting of 1 μg of gDNA, 500 nM primers, 25 μL NEBNext Q5 HotStart HiFi PCR master mix, and nuclease-free water to a final volume of 50 μL. We ran the reactions on a thermal cycler with the following settings: 98°C for 30 seconds, followed by N cycles of 98°C for 10 seconds, then 65°C for 40 seconds, and finally 65°C for 5 minutes. After the PCR, we purified libraries using 35 μL (0.7X) Ampure XP magnetic beads with two 80% ethanol washes followed by final elution in 20 μL 0.1X TE (1 mM Tris HCl pH 8.0 100 μM EDTA). Purified libraries from the same sample were pooled and quantified using the Qubit dsDNA High Sensitivity assay (ThermoFisher), then sequenced on a NextSeq 500 using 150 cycles for read 1 and 8 cycles for each index. We recovered barcodes from sequencing data using custom Python scripts available at: https://github.com/arjunrajlaboratory/timemachine. These scripts search through each read to identify sequences complementary to our library preparation primers, and if these sequences pass a minimum length and phred score cutoff, then the intervening barcode sequence is counted. In addition to counting total reads for each barcode, we also counted the number of "UMIs" incorporated into the library preparation primers and normalized the actual barcode count by the UMI count prior to setting a cutoff for the experiment that applied to all samples in that experiment in order to account for slight variations in sequencing depth between samples. Using the STARCODE software 36 (available at https://github.com/gui11aume/starcode), we merged highly similar barcode sequences (Levenshtein distance ≤ 8), summing the counts and keeping only the more abundant barcode sequence.

### Single-cell RNA sequencing

We used the 10X Genomics scRNA-seq kit v3 (PN-1000121, PN-1000128, PN-1000213) to sequence barcoded cells. We resuspended the cells (targeting ∼10,000 cells for recovery per sample) in PBS and followed the protocol for the Chromium Next GEM Single Cell 3′ Reagent Kits v3.1 as per manufacturer directions (10X Genomics). In brief, we generated gel beads-in-emulsion (GEMs) using the 10X Chromium system and subsequently extracted and amplified (11 cycles) barcoded cDNA as per post-GEM RT-cleanup instructions. We then used a fraction of this amplified cDNA (25%) and proceeded with fragmentation, end-repair, poly A-tailing, adapter ligation, and 10X sample indexing per the manufacturer’s protocol. We quantified libraries using the High Sensitivity dsDNA kit (Thermo Fisher Q32854) on Qubit 2.0 Fluorometer (Thermo Fisher Q32866) and Bioanalyzer 2100 (Agilent G2939BA) analysis prior to sequencing on a NextSeq 500 machine (Illumina) using 28 cycles for read 1, 55 cycles for read 2, and 8 cycles for i7 index.

### Barcode recovery from scRNA-seq data

Since the barcodes are transcribed, we extracted the barcode information from the amplified cDNA from 10X Genomics V3 chemistry protocol (step 2). We ran a PCR side reaction with one primer that targets the 3′ UTR of GFP and the other that targets a region introduced by the amplification step within the V3 chemistry of 10X genomics (read 1). The two primers amplify both the 10X cell-identifying sequence as well as the 100 bp barcode. The number of cycles, typically between 12–15, is decided by the Ct value from a quantitative PCR reaction (New England Biolabs M0543) for the specified cDNA concentration with the following thermal cycler settings: 98 °C for 30 s, *N* cycles of 98 °C for 10 s, 65 °C for 2 min and, finally, 65 °C for 5 min. Following PCR reaction, we immediately performed a 0.7× bead purification (Beckman Coulter B23318) followed by final elution in nuclease-free water. Purified libraries were quantified with High Sensitivity dsDNA kit (Thermo Fisher) on Qubit Fluorometer (Thermo Fisher), pooled, and sequenced on a NextSeq500. We sequence 26 cycles on read 1 which gives 10X cell-identifying sequence and UMI, 124 cycles for read 2, which gives the barcode sequence, and 8 cycles for index i7 to demultiplex pooled samples. The primers used are provided in Supplemental Table 2.

### Computational analyses of scRNA-seq expression data

We adapted the cellranger v3.0.2 by 10X Genomics into our custom pipeline to map and align the reads from NextSeq sequencing run(s) as in^20^. In brief, we downloaded bcl counts and used cellranger mkfastq to demultiplex raw base call files into library-specific FASTQ files. We aligned the FASTQ files to the hg19 human reference genome and extracted gene expression count matrices using cellranger count while also filtering and correcting cell identifiers and unique molecular identifiers (UMI) with default settings. We then performed the downstream single-cell expression analysis in Seurat v3. Within each experimental sample, we removed genes that were present in less than three cells, as well as cells with less than or equal to 200 genes. We also filtered for mitochondrial gene fraction, which was dependent on the cell type. For non-identically treated samples, we integrated them using Harmony. We used these integrated datasets to generate data dimensionality reductions by PCA and UMAP, using 50 principal components for UMAP generation. We tested a range of resolutions with Seurat’s FindClusters command.

### Computational analyses of barcoded single-cell datasets

The barcodes from the side reaction of single-cell cDNA libraries were recovered by developing custom shell, R, and Python scripts (see data availability). In brief, we scan through each read, searching for sequences complementary to the side reaction library preparation primers, filtering out reads that lack the GFP barcode sequence, have too many repeated nucleotides, or do not meet a phred score cut-off. Since small differences in otherwise identical barcodes can be introduced due to sequencing and/or PCR errors, we merged highly similar barcode sequences using STARCODE software, available at https://github.com/gui11aume/starcode. For varying lengths of barcodes (30, 40 or 50, see the pipeline guide provided) depending on the initial distribution of Levenshtein distance of non-merged barcodes, we merged sequences with Levenshtein distance ≤8, summed the counts, and kept only the most abundant barcode sequence. The decision to use a Levenshtein distance ≤8 was reached by systematically analyzing the difference between the experimentally observed mean Levenshtein distance with the theoretically provided mean Levenshtein distance for a pair of barcodes. Previously comparisons of Levenshtein distances for these barcodes found that a Levenshtein distance ≤8 resulted in the least difference between observed and expected mean distances between barcodes ^20^. For the next processing steps and downstream analysis, we first filtered out all barcodes that were associated below the minimum cut-off (dependent on sequencing depth) of unique molecular identifiers (UMI). We next removed all barcodes where one 10X cell-identifying sequence was associated with more than one unique barcode. This could either result from multiplets introduced within gel beads-in-emulsions or because of the same cell receiving multiple barcodes during lentiviral transduction. After these two filtering steps, the qualifying barcodes were used to complete the downstream clone-resolved analyses in which we pulled the UMAP embeddings to illustrate where each cell with a specific sequenced barcode falls on the UMAP and to compare the cluster distributions for each barcode between experiment conditions.

### Bulk RNA sequencing

We sequenced mRNA from WM989-A6 melanoma cells. Each sample replicate came from a separate passage and was processed separately. We used the Single Cell/Low Input RNA Library Prep Kit for Illumina. We sequenced each sample at a depth of approximately 20 million reads on a NextSeq2000 (50 base pair length). We pseudoaligned our reads to hg19 using Kallisto and imported the Kallisto output into R using tximport. The edgeR package was used to normalize reads using the trimmed mean of M values (TMM) method and to calculate counts per million (CPM) per gene. To identify potential learned genes, we first compared the DMSO (14d) and the dexamethasone (14d) samples to and selected genes that were upregulated by dexamethasone treatment (Log2FC threshold of 3). We then selected from this list of genes the ones whose expression decreased from that of the dexamethasone (14d) treatment if dexamethasone was removed (Log2FC threshold of 2). From this list of genes, we then filtered out genes that were massively turned on by trametinib alone by comparing the trametinib condition to the DMSO (14d) condition (Log2FC threshold of 5). Then, to identify genes that may have been learned, we selected genes from this list whose expression increased by a Log2FC of 1 in the dexamethasone to trametinib conditions versus the DMSO to trametinib condition. Finally, with this list of genes, we conducted a literature search to confirm genes that had been shown to be turned on by treatment of dexamethasone or affected by glucocorticoid signaling. We selected 6 genes from this list to create FISH probes to target and selected the first 3 probes that validated (spots seen with dexamethasone treatment)-those being *FKBP5*, *LTBP1*, and *TGM2*.

### ATAC sequencing and analysis

We performed ATAC sequencing according to^30^. Briefly, we lysed the cells and set up the transposition reaction with the Tn5 transposase (generously provided by our collaborators in the Kavitha Sarma lab at The Wistar Institute) at 37℃ for 30 minutes. We isolated the transposed and adaptor-ligated genomic DNA from the reaction with the Zymo DNA Clean and Concentrator--5 Kit (Zymo #D4014) and then amplified the libraries using the custom primers described in Supplemental Table 4 (originally designed by the William Greenleaf lab at Stanford University). We sequenced our libraries on a NextSeq2000 with a P3 100-cycle kit (illumina 20040559), with 60 bp paired-end reads at a depth of approximately 25 million reads per sample.

We aligned our reads to the hg19 assembly with bowtie2 (version 2.8.0) and called peaks using MACS2 (version 2.2.9.1). We then filtered peaks to only those found in at least two samples and for peaks that were not listed in ENCODE blacklist. Using these peak files, we then performed differential peak analysis using DESeq2 (version 1.42.1). We next performed motif analysis on our set of differential peaks using chromVAR (version 1.24.0)^36^, the JASPAR2020 library of transcription factor motifs and motifmatchR package (version 1.24.0) from bioconductor. The default chromVAR code was modified to create expectations only as the average of the 3 days of DMSO condition as opposed to the average of all conditions. Two separate sets of peaks were provided as input into the chromVAR workflow. First was the set of all peaks identified using DESeq2. Second was the set of dexamethasone peaks defined as those that had a fold change greater than 2 in the 3 days of dexamethasone case compared to 3 days of DMSO case. Those dexamethasone peaks were further filtered for peaks not higher in the 3 days of DMSO to 11 days of trametinib condition compared to 3 days of DMSO.

## Data and code availability

Raw data and code are accessible here (https://www.dropbox.com/scl/fo/mx4ef9lvs3bzw8ms18jwt/ABv0e0o_njpT0ooPzNNbgC8?rlkey=0vhml03taajkdkoma0azg4k5t&dl=0).

