## supplemental information for "AP-1 Mediates Cellular Adaptation and Memory Formation During Therapy Resistance"

#### Supplemental Figures and Figure Legends

### Supplemental Figure 1

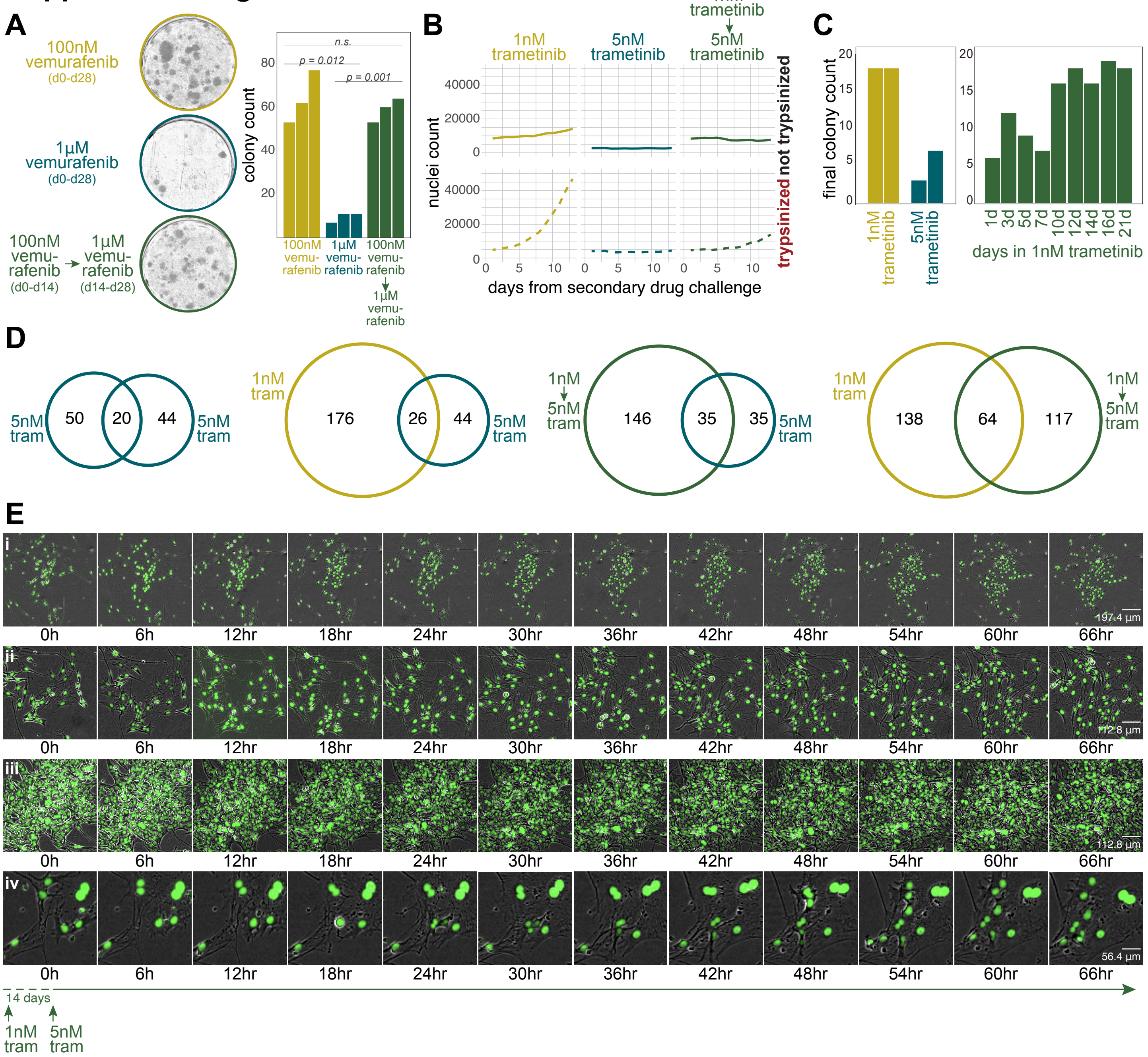

**Supplemental Figure 1. Additional evidence for cellular adaptation during acquisition of therapy resistance**

**(A)** Vemurafenib dose-escalation experiment: Representative images of WM989 A6-G3 colonies at day 28 for low dose alone, high dose alone, and low-to-high-dose switch conditions. Bar plots show average resistant colony counts from 3 biological replicates. Error bars represent standard error of the mean (SEM). Statistical analysis: one-way ANOVA with Shapiro-Wilk's test for normality, Levene's test for variance homogeneity, and post-hoc comparisons using paired t-tests.

**(B)** Test of resistant colony density on trametinib dose-escalation response. Graph shows nuclei count over time, with day 0 representing the dose switch/re-plating day.

**(C)** Additional time course of low-to-high-dose switch (24-well format): Left: Bar plots comparing resistant colony numbers under continuous low versus high dose conditions (two technical replicates each). Right: Bar plots showing resistant colony numbers when switching to high dose after varying low dose exposure durations.

**(D)** Venn diagrams of barcode overlap in DNA barcode lineage tracing experiment for low dose, high-dose alone, and low-to-high-dose switch conditions. (left to right) high dose condition compared with a technical replicate of high dose condition from the same experiment, low dose condition compared with high dose condition, low-to-high-dose switch condition compared with high dose condition, low dose condition compared with low-to-high-dose switch condition.

**(E)** High-resolution time-lapse imaging of colonies immediately following the switch from low to high dose. The top row shows images of the full colony from Figure 2F, while the lower panels show examples of three additional colonies. All colonies shown here developed in the presence of 1nM trametinib, with day 0 marking the start of the switch into high dose.

Since much of the initial characterization of this system[11] was done with vemurafenib, we first confirmed that the phenomenon of adaptation seen with trametinib is also replicated with BRAF$^{V600E}$ inhibitor vemurafenib. As in the trametinib data, treatment with low-dose vemurafenib (100nM) for 14 days prior to challenge with high-dose vemurafenib (1µM) allowed for the survival of colonies that otherwise would not have survived in high-dose vemurafenib (1µM). These data demonstrated that the phenomenon of adaptation also occurs with vemurafenib treatment **(Supplemental Figure 1A).** Next, to rule out the possibility that the increased ability of cells to survive high-dose trametinib (5nM) is due to the increased cell density within the small colonies that formed by the time of the switch to high-dose trametinib, we compared the usual protocol of growing the cells in one well for the duration of the experiment to a protocol in which we trypsinized cells on the day of dose-switch and re-plated the same number of cells as were plated at the start of the experiment. As expected, we found that in splitting and re-plating cells, there was more notable growth by cell count as compared to growing the cells in one well throughout, presumably due to decreased contact inhibition. Additionally, amongst the trypsinized conditions, cells grown in low-dose for the first 14 days grew at a higher rate when replated and exposed to high-dose trametinib as compared to cells that have survived high-dose and then replated and exposed to the same high-dose trametinib. These data suggest that cell density does not protect cells during the low-to-high-dose switch **(Supplemental Figure 1B)**. As an additional control, we repeated the time course from Figure 1C in a 24-well plate format, and we still saw that it took about 12-16 days of low-dose treatment for the number of surviving colonies to plateau. These data provide further support that adaptation is time-dependent **(Supplemental Figure 1C)**.

In addition to quantifying the number of DNA lineage barcodes that survived across the various conditions from Fig. 2D. We also analyzed the overlap of specific barcodes across conditions; our analysis revealed barcodes that survived specifically in the low-to-high-dose switch condition but not in the high dose alone, suggesting that these cells adapted to the therapy during low dose exposure. We also observed that twins of cells surviving the low-to-high-dose switch condition were also likely to survive in the low-dose alone condition

**(Supplemental Figure 1D)**. This finding supports the conclusion that the additional cells surviving at low dose are the ones that persist through the low-to-high-dose switch, rather than a subset of cells inherently resistant to high dose treatment.

As a complement to the time-lapse images in Figure 2F, high-resolution time-lapse imaging in **Supplemental Figure 1E** shows colonies during the switch from low to high dose (upper panels: full colony from Figure 2F, lower panel: another example colony). These time-lapse images of two representative colonies demonstrated several key points: 1. No notable cell death was occurring, which we would have expected to see if progressive selection was occurring within these colonies. 2. The images showed clear cell proliferation during the high-dose challenge. These observations rule out the possibility of further progressive selection within low-dose surviving colonies when subsequently challenged with high-dose trametinib.

### Supplemental Figure 2

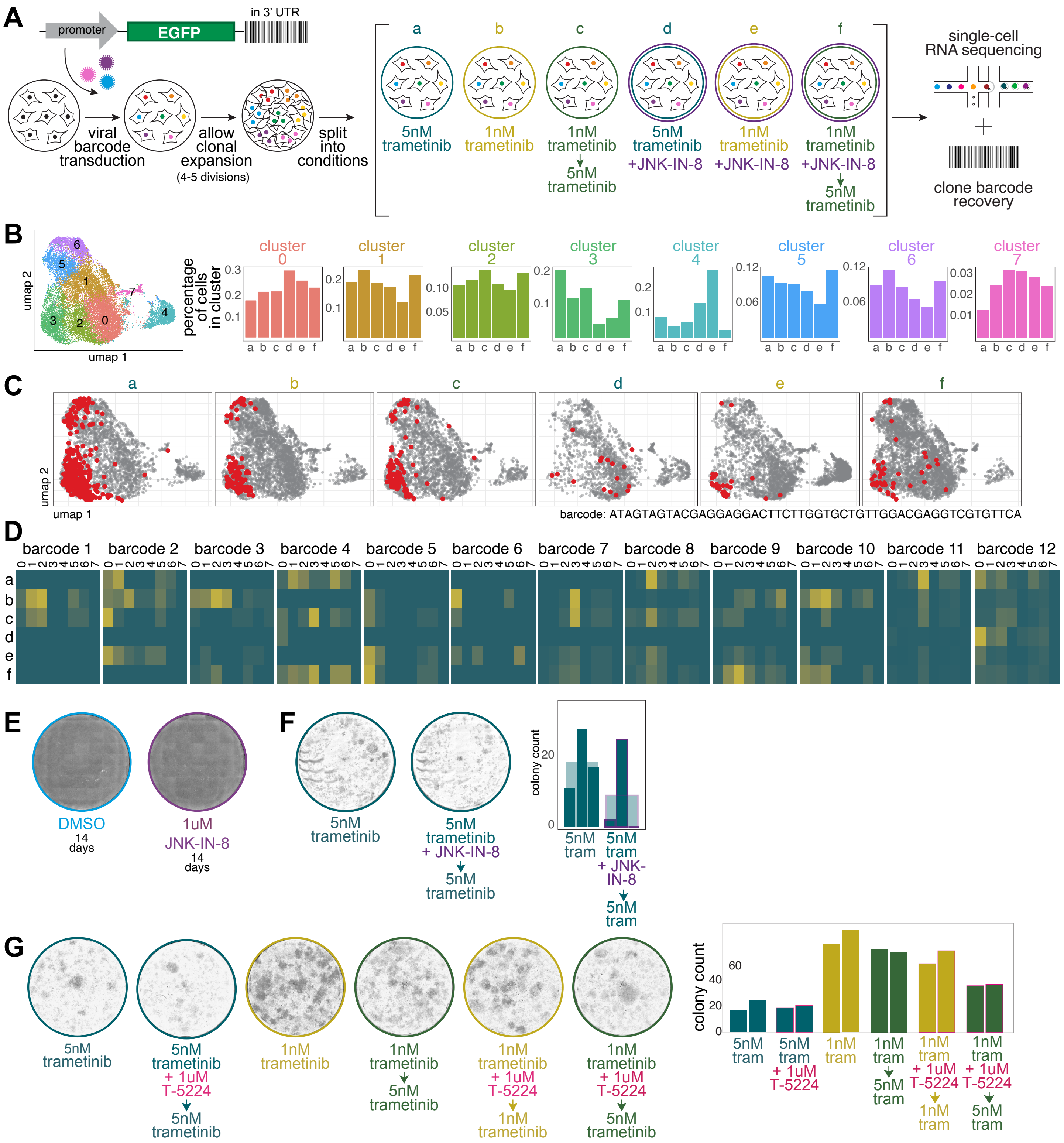

**Supplemental Figure 2: Extended evidence shows cellular adaptation in therapy resistance is dependent on AP-1**

**(A)** Schematic illustrating FateMap experiment assessing transcriptional heterogeneity across resistant lineages under different treatment conditions: WM989 A6-G3 cells were transduced with the FateMap barcode library (MOI ≈ 0.2). After 4–5 cell divisions, the barcoded population was sorted and split into illustrated conditions. Treated cells underwent single cell RNA-seq and barcode sequencing (Table 4.3). Samples are designated as conditions a-f in subsequent panels.

**(B)** Left: UMAP showing cluster identities from all samples integrated using Harmony and clustered with Seurat. Right: Cluster ratio analysis depicting the percentage of cells from each condition in each cluster.

**(C)** UMAPs displaying the transcriptional states of representative twin clones (cells sharing the same barcode) across the six experimental conditions. The unique barcode sequence for this example is shown in the bottom right corner of each set of UMAPs.

**(D)** Heatmap of 12 barcodes (lineages) in 12 separate blocks. Rows represent experimental conditions; columns represent clusters from panel G.

**(E)** Effect of JNK-IN-8 alone on the growth of therapy-naive cells after 14 days of treatment with either DMSO or 1μM JNK-IN-8 (n=1).

**(F)** Effect of the 5 nM trametinib treatment alone compared to the effect of co-treatment of 5 nM trametinib with 1μM JNK-IN-8. Left: Representative whole-well images. Right: Bar plot showing colony counts for 3 biological replicates

**(G)** Effect of AP-1 inhibition using T-5224. Left: Representative whole-well images at day 14 (midpoint and time of dose-switch) and at day 28 (endpoint) showing colony formation under different treatment conditions. Right: Bar plot quantifying final colony counts for each condition. Each bar represents a biological replicate (n=2).

To complement the DNA lineage barcoding shown in Figure 2D, we combined lineage barcoding with single-cell RNA sequencing. This approach allowed us to compare transcriptional state changes in resistant colonies that emerged under the various treatment conditions described **(Supplemental Figure 2A)**. We integrated the reads from scRNAseq with Harmony. We visualized those clusters on a UMAP to ensure that colonies that survived in the low-to-high-dose switch condition were not in a completely new cell state that was not present in the low-dose or high-dose alone conditions. When we separated out each cluster identified via Seurat and compared, for each experimental condition, the percentage of the cells recovered in that condition fell into that cluster **(Supplemental Figure 2B).** Our analysis revealed no significant changes in cluster composition across conditions and no condition-specific clusters, suggesting that the transition from low to high-dose treatment does not induce entirely new cellular fates beyond those observed in either low or high-dose conditions alone. To examine the fate of twin clones (cells sharing the same barcode) across different conditions, we traced the locations of cells with a specific barcode on the UMAP for each condition. In this example, we observed that cells with the selected barcode showed similar cluster distributions in the low dose, low-to-high-dose switch, and high dose conditions. Notably, cells with this barcode from conditions treated with JNK-IN-8 appeared to be distributed across clusters and areas where these barcoded cells were not found in conditions without JNK-IN-8 treatment. In summary, twin clones largely maintained similar transcriptional fate types across conditions but showed increased transcriptional heterogeneity (more cells of the same barcode appearing in different clusters) when co-treated with JNK-IN-8 **(Supplemental Figure 2C)**. We then sampled 11 additional barcodes from the FateMap dataset. We present heatmaps of these 12 barcodes (BC1-BC12) in 12 separate blocks with. The heatmaps reveal that for the majority of barcodes (8/12), the cell fate distribution in the low-to-high-dose switch condition closely resembles that of the low-dose conditions. For some barcodes (2/12), the low-to-high switch condition shows a distribution more similar to the high-dose condition. One barcode (1/12) doesn't completely match either the low dose or high dose patterns, while another (1/12) shows a similar distribution across all conditions **(Supplemental Figure 2D)**. Importantly, we observed that co-treatment with JNK-IN-8 seemed to lead to more spread across clusters for each barcode lineage. These data suggest that cells largely maintain their transcriptional state when transitioning from low to high dose, but this persistence of cell fate can be altered by JNK-IN-8 treatment.

A crucial control for the JNK inhibitor experiments was to verify that treatment with JNK-IN-8 alone at the 1μM dose does not inherently cause cell death or prevent cell growth in the WM989 melanoma system. We observed cells treated with 14 days of JNK-IN-8 (1μM) grow to be confluent in that time **(Supplemental Figure 2E)**. This result indicates that the duration and dose of JNK-IN-8 used in these experiments do not cause drastic cell death or prevent cell growth, showing that any observed effects are not due to JNK-IN-8 toxicity alone.

We also wondered how much adaptation is necessary for the initial transition from primed to resistant. We test this by co-treating naive cells with high dose trametinib and 1μM JNK-IN-8 for the first two weeks before we remove the JNK-IN-8 and allow cells to continue growing 5 nM trametinib **(Supplemental Figure 2E)**. We observe a general decrease in the number of resistant colonies formed in this condition as compared to treatment with high dose trametinib alone for the entire 4 weeks. This supports the notion that cellular adaptation is important for generally surviving this dose (i.e. simply being primed at the point of therapy initiation is not enough to become resistant if JNK is severely inhibited).

In addition to inhibiting AP-1 through JNK inhibition, we sought to replicate the results with another pharmacological inhibitor of AP-1, T-5224, a small molecule inhibitor that blocks the DNA binding activity of c-Fos/c-Jun **(Supplemental Figure 2G)**. In two biological replicates, we observed that co-treatment of 1μM T-5224 with low dose trametinib (1nM) for two weeks followed by treatment with low dose trametinib alone does not drastically alter the number of colonies that survived and became resistant at the end of the experiment compared to cells that only received low dose for that duration. To note, with this dose and treatment duration of T-5224, there is not a notable effect on the survival of cells exposed to high dose of trametinib. However, for cells that received co-treatment of 1μM T-5224 with low dose trametinib (1nM) for two weeks followed by treatment with high dose trametinib (5nM), there was a decrease in the number of final colonies. These data demonstrate that a second pharmacological inhibitor of AP-1 replicated the effects seen with JNK inhibition, further supporting the role of AP-1 in cellular adaptation to therapy.

### Supplemental Figure 3

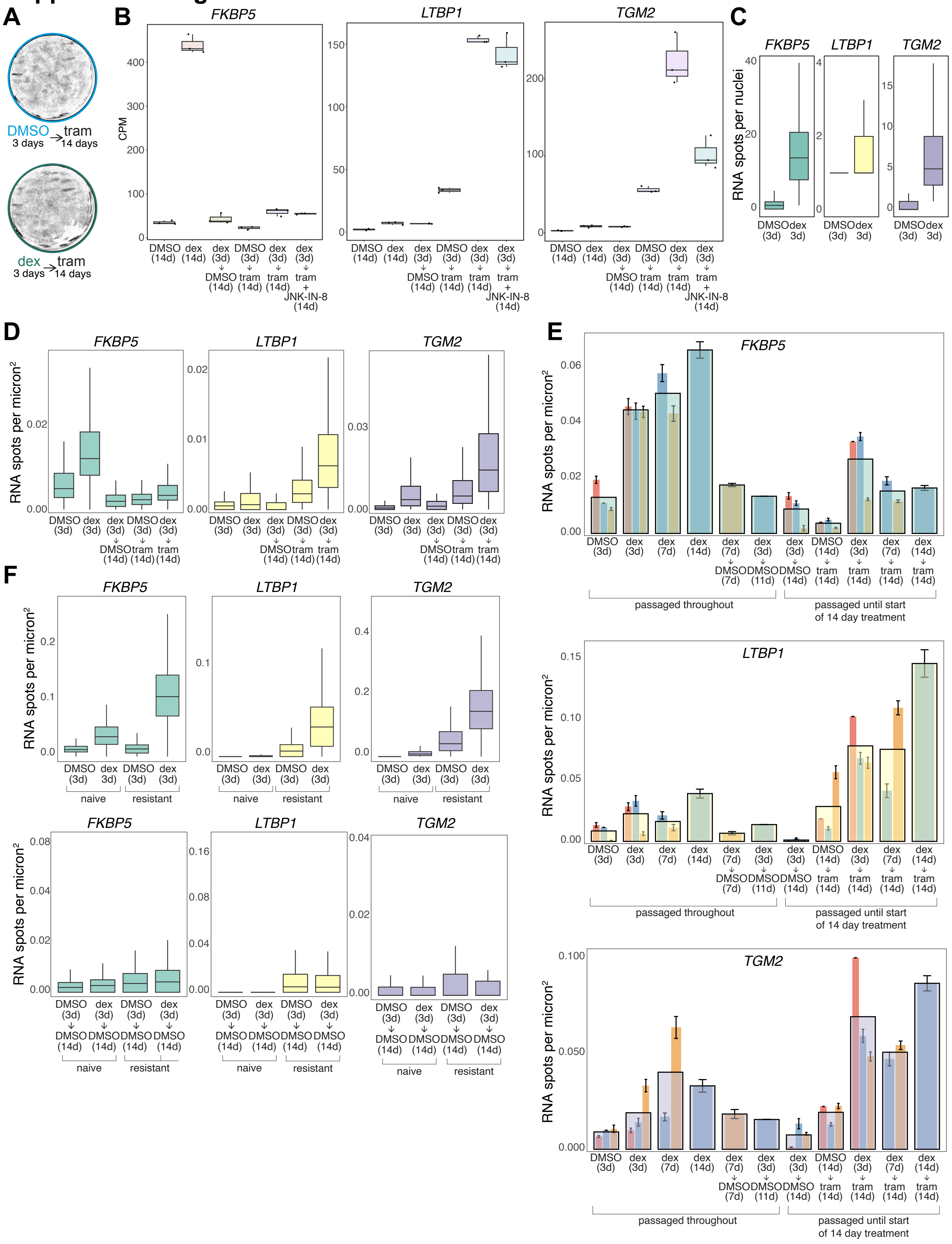

**Supplemental Figure 3. Validation of dexamethasone-induction and reversion of gene expression in naive and resistant cells**

**(A)** Phase-contrast images of WM989-A6-G3 cells grown in various conditions
**(B)** Bulk RNA sequencing data for *FKBP5*, *LTBP1*, and *TGM2* under various conditions as shown.
**(C)** Boxplots showing gene expression (RNA counts per nucleus) in crowded cell conditions (n=1).
**(D)** Single-molecule RNA FISH quantification of *FKBP5*, *LTBP1*, and *TGM2,* expression under various conditions in a different clonal population of WM989 cells (H2B-GFP WM989 A6-G3-A10)
**(E)** Single-molecule RNA FISH quantification of *FKBP5*, *LTBP1*, and *TGM2* expression under various dexamethasone treatment durations. Data from 3 biological replicates (not all conditions present in each replicate). Larger overlaying bar plots represent the average of each condition. Smaller underlying bars represent individual replicate averages, consistently color-coded across conditions and genes. Error bars show standard error of the mean (SEM). The number of days spent in each condition are noted below each treatment.
**(F)** Comparison of dexamethasone response and memory in naive and resistant WM989 cells. (left plots) boxplots comparing the induction of *FKBP5*, *LTBP1*, and *TGM2* expression in naive versus resistant cells in response to dexamethasone treatment. (right plots) Boxplots comparing the decrease in expression of *FKBP5*, *LTBP1*, and *TGM2* in naive versus resistant cells after dexamethasone removal.

To demonstrate that the transition from primed to fully resistant cells induces the formation of cellular memories, we tested whether we could cause the formation of a memory of a "passenger" gene's expression in response to dexamethasone as an inducer. We first tested to make sure that 3 days of dexamethasone treatment (the primary exposure duration we would use), would not change the ability of these cells to survive and form colonies if treated with 1 nM trametinib **(Supplemental Figure 3A)**. We observed no obvious changes in colony formation in 1 nM trametinib after 3 days of dexamethasone treatment compared to 3 days of DMSO treatment.

To identify potential evidence of memory formation, we conducted bulk RNA sequencing on WM989 cells under various treatment conditions as shown **(Supplemental Figure 3B)**. We found a subset of genes that showed potential for learning, designed single-molecule RNA FISH probes for six of these genes, and proceeded with the first three probe sets to test our hypothesis (*FKBP5*, *LTBP1*, and *TGM2*). These genes displayed significantly higher induction when dexamethasone treatment was followed by trametinib, compared to trametinib treatment alone, and this enhanced expression was reduced when JNK-IN-8 was added, suggesting that learning occurs at loci responsive to trametinib and is partially dependent on AP-1 activity.

After seeing the results from the experiment in Figure 3B, we conducted a series of experiments to rule out possible confounding factors that may suggest alternative explanations to the memory formation we observed. Since we had observed in the single-molecule RNA FISH data collected in Figure 3B that conditions where cells were grown in dexamethasone for 3 days followed by DMSO for 14 days showed a global downregulation of gene expression, including in the expression of housekeeping gene hUBC, we induced target gene expression with 3 day dexamethasone treatment in high cell density conditions **(Supplemental Figure 3C)**. The induction of dexamethasone target genes at high cell density ruled out the possibility that the observed return to baseline expression in our main experiments was an artifact of high cell density.

We also performed the dexamethasone-induced passenger memory experiment in a different clonal cell line of WM989s to further validate our finding from Figure 3B **(Supplemental Figure 3D)**. The results here closely resemble the results seen within Figure 3B, suggesting that the findings from Figure 3B are not an artifact of that single clonal population. (Note that JNK inhibition was not tested in this experiment with the H2B-GFP

WM989 A6-G3-A10 cells.

Since bulk RNA sequencing and target gene selection were based on 14 days of dexamethasone exposure while the validation by smFISH was based on 3 days of dexamethasone exposure, we measured the effects of a longer dexamethasone exposure on the expression of *FKBP5*, *LTBP1*, and *TGM2* **(Supplemental Figure 3E)**. We found a general trend of increased gene expression as the duration of dexamethasone exposure increased and a decrease in gene expression towards baseline when dexamethasone was removed, demonstrating that longer dexamethasone treatment generally led to higher gene induction. Importantly, in this experiment, cells were passaged every 3-4 days (n=3; not all conditions were represented in all replicates). In these conditions, gene expression increased during treatment and decreased upon dexamethasone removal despite cell passaging, demonstrating the reversibility of the effect and ruling out potential changes in expression resulting from prolonged culture without passaging. We also used this time course to test trametinib-induced memory to see the dynamics of memory with a longer dexamethasone exposure time while also passaging the cells prior to exposure to trametinib to prevent overcrowding. We observed a general trend where we observed a stronger trametinib-induced memory effect for *FKBP5*, *LTBP1*, and *TGM2* by single-molecule RNA FISH. Importantly, these results demonstrate that passaging cells during the experiment does not alter the memory formation and retention effects we have observed. This control supports the robustness of our findings across different experimental conditions and timescales, reinforcing the validity of our cellular memory model in the context of drug resistance acquisition.

To eliminate the possibility that resistant cells have different induction and reversion timescales compared to therapy-naive cells, we conducted an experiment using a population of resistant cells that had never been exposed to dexamethasone. Resistant cells (and, as a control, naive cells) were treated with dexamethasone for 3 days, followed by 11 days of culture without dexamethasone **(Supplemental Figure 3F)**. We observed that the expression of our target genes (*FKBP5*, *TGM2*, and *LTBP1*) increased upon dexamethasone induction and reverted to baseline levels after dexamethasone removal. This control experiment (n=1) showed that resistant cells do not have an inherently altered regulatory system for these genes, as they exhibit similar patterns of gene induction and reversion to baseline as naive cells. In line with our other experimental observations, this evidence points to a critical time window for memory formation that occurs early in the trametinib treatment process.

### Supplemental Figure 4

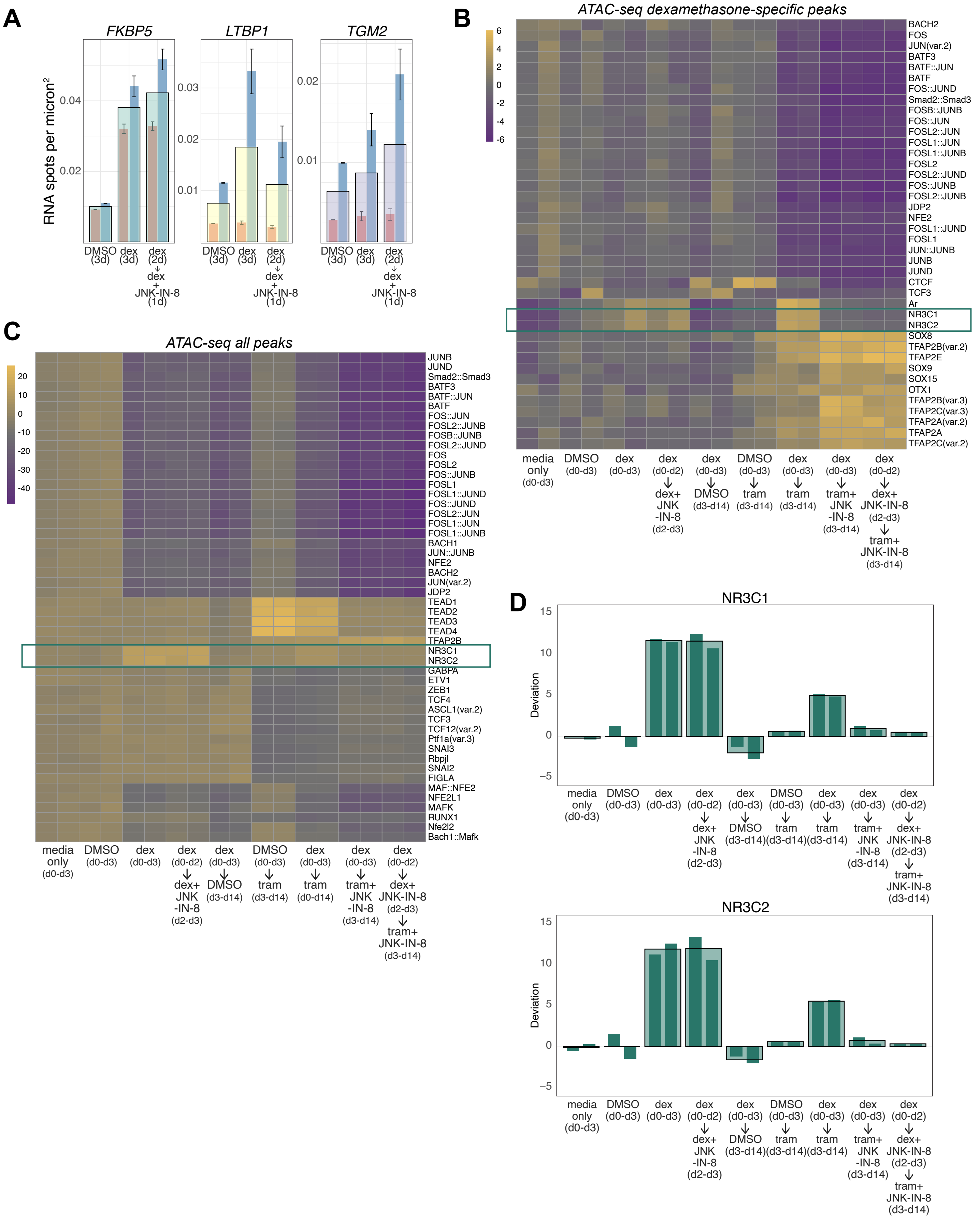

**Supplemental Figure 4. Validation of AP-1 inhibition by JNK inhibition and comprehensive chromatin accessibility analysis support an AP-1 dependent mechanism of memory formation.**

**(A)** Bar plots showing results from two biological replicates of the effects of dexamethasone treatment for two days followed by co-treatment of JNK-IN-8 (1μM) and dexamethasone for one day. Larger overlaying bars represent the average of each condition. Smaller underlying bars show individual replicate averages, consistently color-coded across conditions and genes. Error bars indicate SEM.
**(B)** chromVAR heatmap showing changes in transcription factor signatures in all experimental conditions for peaks with increased chromatin accessibility after 3 days of dexamethasone treatment compared to 3 days of DMSO and peaks that were not induced by trametinib treatment alone. (A subset of this heatmap is shown in Figure 3D).
**(C)** chromVAR heatmap demonstrating changes in Glucocorticoid receptor and AP-1 transcription factor signatures in experimental conditions for all variable peaks.
**(D)** Bar graphs of chromVAR deviation z-scores demonstrating changes in transcription factor signatures for NR3C1(top) and NR3C2 (bottom) for all conditions performed in the experiment. Large bars represent the mean for each condition and smaller underlying bars show the 2 replicates. A subset of this data is shown in Figure 3D.

In Figure 3B, we showed the effects of inhibiting AP-1 with JNK-IN-8 overlapping with the last day of dexamethasone induction. To rule out the possibility that JNK-IN-8 treatment on the final day of a three-day dexamethasone exposure affected the expression of our target genes, we measured expression of *FKBP5*, *LTBP1*, and *TGM2* by single-molecule RNA FISH at the end of the three-day treatment **(Supplemental Figure 4A)**. We found that JNK-IN-8 treatment on the third day of a three-day dexamethasone exposure did not significantly alter the expression of these genes as compared to dexamethasone treatment alone for 3 days, suggesting that the decreased gene expression observed when JNK-IN-8 and trametinib are co-administered in our main experiments is likely due to the inhibition of AP-1 during the memory formation process, rather than an immediate effect of JNK-IN-8 on dexamethasone-induced gene expression.

To further analyze changes in chromatin accessibility associated with memory formation, we generated a subset of peaks to analyze that showed both an increase in chromatin accessibility in the 3 days of dexamethasone conditions (3 days of dexamethasone conditions compared to 3 days of DMSO conditions) but also no increase simply as a result of treatment with trametinib (3 day of DMSO treatment followed by 11 days of trametinib treatment compared to 3 days of DMSO). This subset of peaks allowed us to focus specifically on the effects of dexamethasone-responsive peaks to the exclusion of the effects of trametinib treatment. In this subset, we saw that accessibility increased in regions with the GR element upon dexamethasone treatment and not with trametinib alone, but when dexamethasone was removed and trametinib was added (the memory formation condition), we observed not only maintenance but actually an increase in chromatin accessibility of regions that contain sequences with GR signatures **(Supplemental Figure 4B)**. We observed similar effects when examining all peaks as we had when comparing peaks specifically responsive to dexamethasone and excluding trametinib-induced peaks **(Supplemental Figure 4C)**. In short, whether we look at all variable peaks or the subset of peaks that are responsible to dexamethasone but not induced by trametinib alone, we see memory of dexamethasone-induced increased chromatin accessibility at GR binding site that is induced by trametinib and blocked by JNK-inhibitor, regardless of whether the JNK inhibition started on the last day of dexamethasone treatment or at the same time as the trametinib treatment. Collectively these results collectively support our findings that AP-1 plays a crucial role in maintaining the chromatin accessibility changes associated with memory formation.

We also extended our analysis to include all experimental conditions as shown in Figure 3D. Bar graphs of chromVAR deviation z-scores demonstrating changes in transcription factor signatures for NR3C1 (top) and NR3C2 (bottom) are presented for all conditions performed in the experiment **(Supplemental Figure 4D)**. Large bars represent the mean for each condition, while smaller underlying bars show the two replicates. This comprehensive analysis revealed the same pattern of changes observed in the heatmaps; specifically, we observed increased accessibility at GR binding sites with dexamethasone treatment, which was maintained or even enhanced when followed by trametinib treatment. Importantly, this maintenance of accessibility was blocked by JNK inhibition, regardless of whether JNK inhibition started on the last day of dexamethasone treatment or concurrently with trametinib treatment. These results further support our conclusion that AP-1 activity is crucial for maintaining the chromatin accessibility changes associated with memory formation.

### Supplemental Figure 5

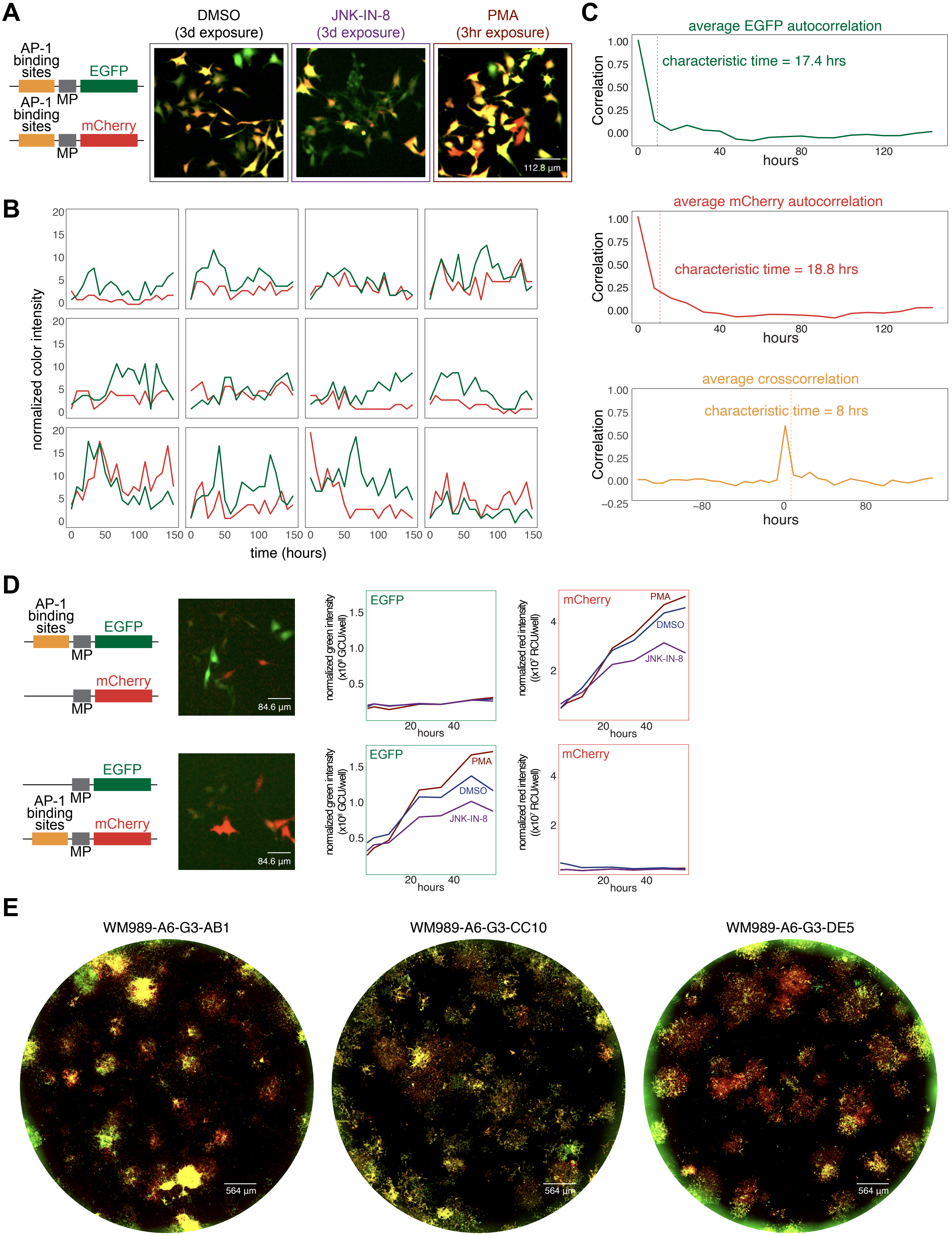

**Supplemental Figure 5. Validation of the dynamics, stability, and specificity of AP-1 reporter cells**
**(A)** Sample images of a single-cell cloned population of WM989 A6-G3 cells transduced with the two AP-1 reporter constructs (WM989 A6-G3-BE5) (left). The right panel shows images collected at 7 days after either a 3-day exposure of DMSO, a 3-day exposure of 1μM JNK-IN-8, or a 3-hour exposure of PMA. All three images are contrasted the same.
**(B)** Twelve independent cell traces of EGFP and mCherry fluorescence intensities. Each graph represents a single WM989A6-G3-BE5 AP-1 reporter cell tracked over 144 hours. Color intensity is calculated as the mean fluorescence intensity of the cell minus the median of the fluorescence intensity of a 10-pixel annulus around that cell. These cell traces show that there are various amplitudes of fluctuations in EGFP and in mCherry.
**(C)** Autocorrelation and cross-correlation analysis for EGFP and mCherry signals (n=34 cells). EGFP autocorrelation shows an average characteristic time of 17.4 hours, while mCherry autocorrelation has an average characteristic time of 18.8 hours. The EGFP-mCherry cross-correlation displays an average correlation time of 8 hours. Characteristic time is calculated as when the correlation drops below a threshold of 1/e of its initial value.
**(D)** Top: normal AP1-EGFP reporter construct and an mCherry construct co-transduced with only the minimal promoter and no AP-1 binding site. Bottom: normal AP1-mCherry reporter construct co-transduced with an EGFP construct with only the minimal promoter and no AP-1 binding site. Clonal populations of each combination were isolated by dilution cloning. To the right of the schematic are day 1 images of a clonal population of cells that have the construct on the left and on the right, respectively. Graphs to the right show Incucyte S3 well measurements of normalized green and red fluorescence intensity values for a clonal population of each the left and right constructs.
**(E)** Whole well images of colonies formed after 28 days of treatment with 1 nM trametinib for 3 different clonal populations of the WM989-A6-G3 reporter cells: (left) WM989-A6-G3-AB1, (middle) WM989-A6-G3-CC10, (right) WM989-A6-G3-DE5.

As a first test of the AP-1 dual reporter system, we confirmed that a clonal population of WM989-A6-G3-BE5 reporter cells showed a notable reduction in fluorescence intensity resulting from treatment with JNK-IN-8 and an overall increase in fluorescence for cells treated with a short-term exposure of PMA, demonstrating the responsiveness of the reporter system to AP-1 inhibition and activation, respectively **(Supplemental Figure 5A)**. We then sought to characterize the baseline fluctuations of fluorescent reporters in therapy-naive cells. **Supplemental Figure 5B** shows twelve independent cell traces of EGFP and mCherry fluorescence intensities for a WM989-A6-G3-BE5 AP-1 reporter cell tracked over 144 hours. These cell traces showed that there were various amplitudes of fluctuations in EGFP and mCherry expression levels. Importantly, we observed instances when the mCherry and EGFP intensities change in the same direction (increased or decreased) from one time point to another and instances when the mCherry and EGFP change in opposite directions from one time point to another. To quantify these fluctuations at baseline, we calculated autocorrelation and cross-correlation for EGFP and mCherry signals (n = 34 cells). EGFP autocorrelation showed an average characteristic time of 17.4 hours, while mCherry autocorrelation had an average characteristic time of 18.8 hours. The EGFP-mCherry cross-correlation displayed an average cross-correlation time of 8 hours. Here characteristic time is calculated as when the correlation drops below a threshold of 1/e of its initial value. These values of characteristic times for EGFP and mCherry autocorrelation and crosscorrelation indicate that expression levels of EGFP and mCherry change roughly daily and that EGFP and mCherry expression do not sustain correlation for long periods **(Supplemental Figure 5C)**.

We also sought to verify that the reporter system was specifically dependent on the AP-1 binding sites. To achieve this, we created an internally controlled system where we either co-transduced into WM989 A6-G3 cells the normal AP1-EGFP reporter construct and an mCherry construct with only the minimal promoter and no AP-1 binding sites, or we co-transduced the normal AP1-mCherry reporter construct with an EGFP construct with only the minimal promoter and no AP-1 binding sites **(Supplemental Figure 5D)**. These images confirm that there is very low but present expression of the fluorescent protein without the AP-1 binding sites. Graphs below show that within the same clonal population, the AP-1 responsive changes in fluorescence were only seen for the fluorescent protein that has the AP-1 binding site upstream, confirming the specificity of the reporter system to AP-1 activity **(Supplemental Figure 5D)**.

To demonstrate that the phenomenon we observed in Figure 4C was not only a result of the exact site of integration of the reporter constructs in the BE5 clone, we isolated and generated trametinib-resistant colonies with 3 additional reporter cell lines, all dilution-cloned and expanded from a single cell **(Supplemental Figure 5E)**. For reporter lines AB1, CC10, and DE5, we observed diverse colors of resistant colonies in all three additional cell lines, as we saw with the BE5 line, suggesting that the memory formation process is a general phenomenon independent of the specific integration pattern of the reporter constructs.

### Supplemental Figure 6

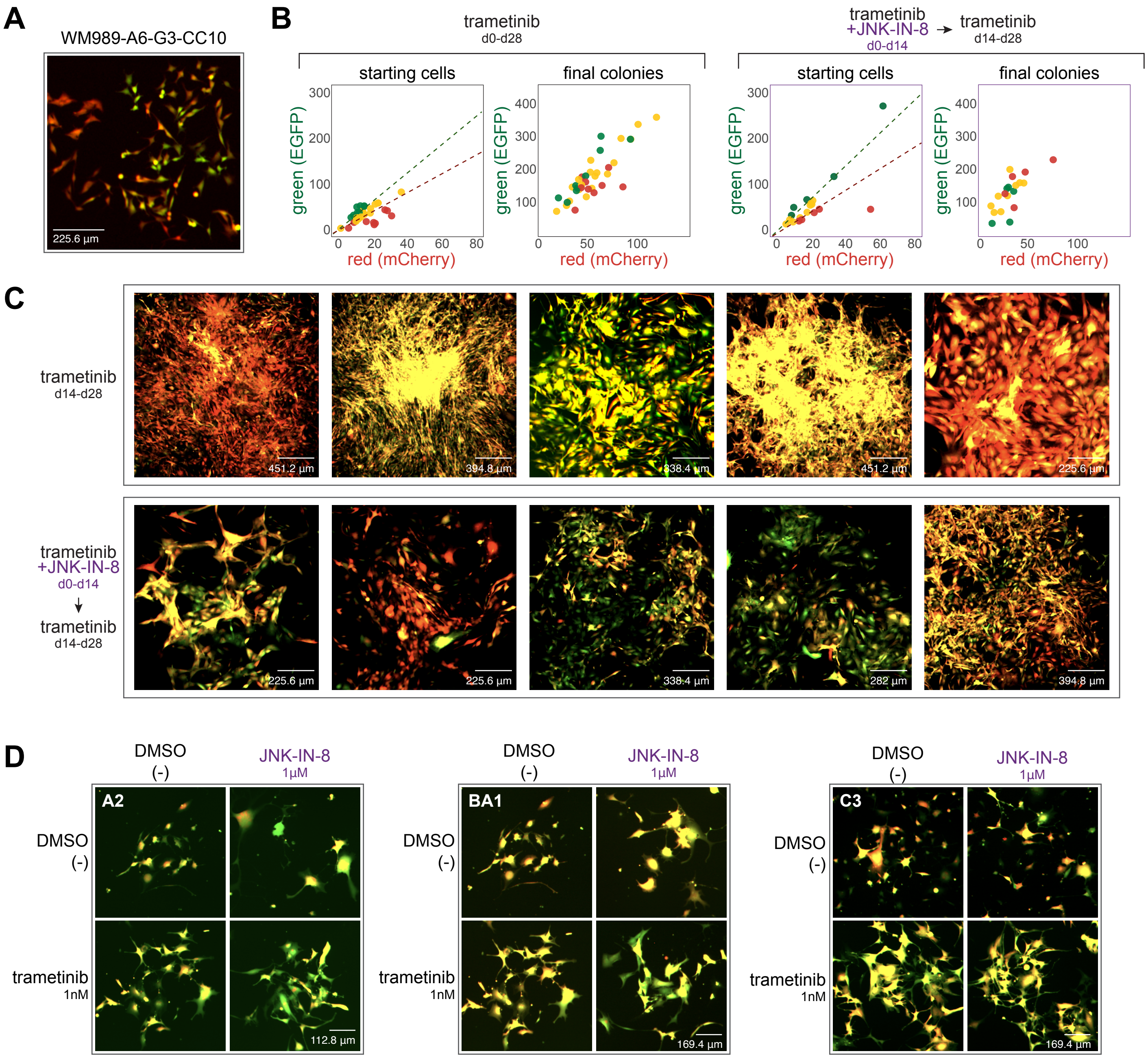

**Supplemental Figure 6. Validation of the dynamics of a second clone of AP-1 reporter cells - WM989-A6-G3-CC10**

**(A)** Image showing cell-to-cell variability in expression in a single-cell-derived clone of WM989 A6-G3 cells transduced with both reporter constructs (WM989-A6-G3-CC10)

**(B)** Scatter plots of fluorescence quantifications from Incucyte S3 images of WM989-A6-G3-CC10 cells grown under two conditions: Left: 1nM trametinib alone for 28 days. Right: 1nM trametinib and 1µM JNK-IN-8 co-treatment for days 0-14, followed by trametinib alone for days 14-28. Colonies were traced back through time-lapse images to identify their originating cells. The initial red-to-green fluorescence intensity ratio of these starting cells was used to categorize the initial 'color' of the cell: top 25th percentile as red, bottom 25th percentile as green, and the remainder as yellow. This initial categorization is used to determine the color coding of the final colonies. The same system is applied for the JNK-IN-8 panel, with the color coding of the final colonies determined according to the starting cell red-to-green ratio for cells in the JNK-IN-8 treatment condition.

**(C)** Day 28 images of the colonies shown in panel A. The top panel shows trametinib alone for the duration of the experiment, while the bottom panel shows trametinib and JNK-IN-8 co-treatment during the first 14 days and trametinib alone for the next 14 days. All images are contrasted the same for each channel.

**(D)** Images of three manually isolated resistant colony clones, A2, BA1, and C3 cultured for more than 1 month after initial colony formation and isolation. Cells were plated at day 0 and either continued to be grown in trametinib or had trametinib removed (DMSO alone), and for both those conditions, with and without 1 µM JNK-IN-8. These images were collected after 7 days of this treatment.

To validate our findings from Figure 4 in another clonal cell line, we first demonstrate the baseline variability of the CC10 clone **Supplemental Figure 6A**. We also performed time-lapse tracking of cells through the process of colony formation in the CC10 clone. We quantified the fluorescence levels of the initial state of the cell and then connected those cells to the ultimate therapy-resistant colony and found that cells that were initially more mCherry or EGFP-positive ended up forming resistant colonies that largely preserved those differences **(Supplemental Figure 6B)**, as we saw for the BE5 clone (Fig. 4E). JNK inhibition led to a reduction in the induction of the levels of both reporters as well as a more noticeable variation of colors within colonies, again suggesting that blocking memory formation allows for cell fates to be more malleable **(Supplemental Figure 6B-C)**. To visualize the durability of therapy resistance memory and its dependence on AP-1, we isolated five trametinib-resistant colonies grown from WM989 A6-G3-CC10 and grew three of these individual therapy-resistant colonies (A2, BA1, C3) with and without trametinib and with and without JNK inhibitor for another 7 days. We found that resistant colonies maintained high expression of both fluorescent proteins after a month in trametinib. This signal persisted largely unchanged when grown with or without trametinib and JNK inhibitor for an additional week, suggesting that once established, the epigenetic memory of resistance is maintained by a mechanism independent of trametinib effects or AP-1 activity **(Supplemental Figure 6D).**

**Tables**

**Table 1: Cloning primers for generation of AP-1 responsive fluorescent reporters**

| primer name | sequence |
|---|---|
| A | attgtcgacctacttgtacagctcgt |
| B | gcgaccggtaagcttcaagatgaggcaa |
| C | atcgatcgatggatcctgagtcagtgactc |
| D | atcgatcgatgaattcagcttgaataaaat |

**Table 2: Primers used to generate gDNA barcode sequencing libraries**

| primer name | sequence | index | stagger |
|---|---|---|---|
| LentiEFS_GFP_NGS_i5.2.1 | AATGATACGGCGACCACCGAGATCTACACTAGATCGCACACTCTTTCCCTACACGACGCTCTTCCGATCTNHNNNNTCGACTAAACGCGCTACTTG | TAGATCGC | None |
| LentiEFS_GFP_NGS_i5.2.2 | AATGATACGGCGACCACCGAGATCTACACCTCTCTATACACTCTTTCCCTACACGACGCTCTTCCGATCTNHNNNNCGTCGACTAAACGCGCTACTTG | CTCTCTAT | CG |
| LentiEFS_GFP_NGS_i5.2.3 | AATGATACGGCGACCACCGAGATCTACACTATCCTCTACACTCTTTCCCTACACGACGCTCTTCCGATCTNHNNNNGACTCGACTAAACGCGCTACTTG | TATCCTCT | GAC |
| LentiEFS_GFP_NGS_i5.2.4 | AATGATACGGCGACCACCGAGATCTACACAGAGTAGAACACTCTTTCCCTACACGACGCTCTTCCGATCTNHNNNNATAGTCTCGACTAAACGCGCTACTTG | AGAGTAGA | ATAGTC |
| LentiEFS_GFP_NGS_i5.2.5 | AATGATACGGCGACCACCGAGATCTACACGTAAGCAGACACTCTTTCCCTACACGACGCTCTTCCGATCTNHNNNNTACGTCGACTAAACGCGCTACTTG | GTAAGCAG | TACG |
| LentiEFS_GFP_NGS_i5.2.6 | AATGATACGGCGACCACCGAGATCTACACACTGCGTAACACTCTTTCCCTACACGACGCTCTTCCGATCTNHNNNNCCTAATCGACTAAACGCGCTACTTG | ACTGCGTA | CCTAA |
| LentiEFS_GFP_NGS_i7.2.1 | CAAGCAGAAGACGGCATACGAGATTAAGGCGAGTGACTGGAGTTCAGACGTGTGCTCTTCCGATCTGTCCTGCTGGAGTTCGTGAC | TAAGGCGA | None |
| LentiEFS_GFP_NGS_i7.2.2 | CAAGCAGAAGACGGCATACGAGATCGTACTAGGTGACTGGAGTTCAGACGTGTGCTCTTCCGATCTAGGTCCTGCTGGAGTTCGTGAC | CGTACTAG | AG |
| LentiEFS_GFP_NGS_i7.2.3 | CAAGCAGAAGACGGCATACGAGATAGGCAGAAGTGACTGGAGTTCAGACGTGTGCTCTTCCGATCTTCAAGTCCTGCTGGAGTTCGTGAC | AGGCAGAA | TCAA |
| LentiEFS_GFP_NGS_i7.2.4 | CAAGCAGAAGACGGCATACGAGATTCCTGAGCGTGACTGGAGTTCAGACGTGTGCTCTTCCGATCTCACTTAAGTCCTGCTGGAGTTCGTGAC | TCCTGAGC | CACTTAA |
| LentiEFS_GFP_NGS_i7.2.5 | CAAGCAGAAGACGGCATACGAGATCGACTCCTGTGACTGGAGTTCAGACGTGTGCTCTTCCGATCTTGTCCTGCTGGAGTTCGTGAC | CGACTCCT | T |

| | | | |
|---|---|---|---|
| LentiEFS_GFP_NGS_i7.2.6 | CAAGCAGAAGACGGCATACGAGATTAGGCATGGTGACTGGAGTTCAGACGTGTGCTCTTCCGATCTCTTGAGTCCTGCTGGAGTTCGTGAC | TAGGCATG | CTTGA |

**Table 3: Sequences of the smFISH probes used in this study**

| gene | sequence | probe |
|---|---|---|
| FKBP5 | cctgtctttttggaggtaa | FKBP5_exonContinuous_1 |
| FKBP5 | cccactcttttgacaatctt | FKBP5_exonContinuous_2 |
| FKBP5 | actttgtctccaatcatcgg | FKBP5_exonContinuous_3 |
| FKBP5 | tccatttgacaattttcctt | FKBP5_exonContinuous_4 |
| FKBP5 | gcctttgccaagactaaaga | FKBP5_exonContinuous_5 |
| FKBP5 | catatctctcctttcttcat | FKBP5_exonContinuous_6 |
| FKBP5 | catattctggtttgcacagt | FKBP5_exonContinuous_7 |
| FKBP5 | gaattttagggagactgcca | FKBP5_exonContinuous_8 |
| FKBP5 | cctttgaaatcaaggagctc | FKBP5_exonContinuous_9 |
| FKBP5 | gtttggttctccggataatg | FKBP5_exonContinuous_10 |
| FKBP5 | tggatttgaatatccctctc | FKBP5_exonContinuous_11 |
| FKBP5 | caggtggatttctactgttg | FKBP5_exonContinuous_12 |
| FKBP5 | attggaatgtcgtggtcttc | FKBP5_exonContinuous_13 |
| FKBP5 | ctccagagctttgtcaattc | FKBP5_exonContinuous_14 |
| FKBP5 | aaccatatcttggtccaaga | FKBP5_exonContinuous_15 |
| FKBP5 | ctcagcattaggttcaatgc | FKBP5_exonContinuous_16 |
| FKBP5 | attctttggccttttcgaag | FKBP5_exonContinuous_17 |
| FKBP5 | gctccaatttttctttggta | FKBP5_exonContinuous_18 |
| FKBP5 | ttctctttgacaatggcagc | FKBP5_exonContinuous_19 |
| FKBP5 | tcccttgaagtatacggttc | FKBP5_exonContinuous_20 |
| FKBP5 | atcttcccatactgaatcac | FKBP5_exonContinuous_21 |
| FKBP5 | tattccatctctaaccagga | FKBP5_exonContinuous_22 |
| FKBP5 | cagaagctttcgattccttt | FKBP5_exonContinuous_23 |
| FKBP5 | cacatggccaggttcagaaa | FKBP5_exonContinuous_24 |
| FKBP5 | ggtgtattctctaagcttca | FKBP5_exonContinuous_25 |
| FKBP5 | ttgtcacagcattcaacagc | FKBP5_exonContinuous_26 |
| FKBP5 | atacaagcctttctcattgg | FKBP5_exonContinuous_27 |
| FKBP5 | ttggctgactcaaactcgtt | FKBP5_exonContinuous_28 |
| FKBP5 | ttttctggcacatggagatc | FKBP5_exonContinuous_29 |
| FKBP5 | gaacatgttggcgtatatcc | FKBP5_exonContinuous_30 |

| | | |
|---|---|---|
| FKBP5 | ttccttttcattagtgacc | FKBP5_exonContinuous_31 |
| FKBP5 | aggtttctcttcttccattg | FKBP5_exonContinuous_32 |
| LTBP1 | tgtctgccttcttagatgaa | LTBP1_exonContinuous_1 |
| LTBP1 | tgagaatgtatctgctgggg | LTBP1_exonContinuous_2 |
| LTBP1 | aatcaccacactctgggaag | LTBP1_exonContinuous_3 |
| LTBP1 | aaagtacttgggcttgagca | LTBP1_exonContinuous_4 |
| LTBP1 | ggccatcaattcttgaaacc | LTBP1_exonContinuous_5 |
| LTBP1 | tgagcttcttttgtcttctg | LTBP1_exonContinuous_6 |
| LTBP1 | ataagatggtttcttggggc | LTBP1_exonContinuous_7 |
| LTBP1 | tacaccttgtagctgacatt | LTBP1_exonContinuous_8 |
| LTBP1 | cccatggtattcaaacactc | LTBP1_exonContinuous_9 |
| LTBP1 | tttgcaggtacatcgatagc | LTBP1_exonContinuous_10 |
| LTBP1 | tgaacagacagagggtgcat | LTBP1_exonContinuous_11 |
| LTBP1 | tcaggagctacttcaacagg | LTBP1_exonContinuous_12 |
| LTBP1 | cactggtaggttaatgcagt | LTBP1_exonContinuous_13 |
| LTBP1 | tcacagtattcacaccggaa | LTBP1_exonContinuous_14 |
| LTBP1 | tggacaagtgcttggattca | LTBP1_exonContinuous_15 |
| LTBP1 | ggcattcatcaatgtcttca | LTBP1_exonContinuous_16 |
| LTBP1 | gatatcttcacagtggtctc | LTBP1_exonContinuous_17 |
| LTBP1 | aaacactcttgtcctccaag | LTBP1_exonContinuous_18 |
| LTBP1 | cagtattaatgcagtctcct | LTBP1_exonContinuous_19 |
| LTBP1 | tgtcacaaaacccgtgactg | LTBP1_exonContinuous_20 |
| LTBP1 | ataacagaggcagcggaagg | LTBP1_exonContinuous_21 |
| LTBP1 | gttcacattcattcacatcc | LTBP1_exonContinuous_22 |
| LTBP1 | attatcacagagactggcgt | LTBP1_exonContinuous_23 |
| LTBP1 | aggggaagatttcgcagtta | LTBP1_exonContinuous_24 |
| LTBP1 | cttgcttacagtagcattca | LTBP1_exonContinuous_25 |
| LTBP1 | aactactggggtcttgacat | LTBP1_exonContinuous_26 |
| LTBP1 | agcatagtcatctgaatcct | LTBP1_exonContinuous_27 |
| LTBP1 | accattttcacatccattga | LTBP1_exonContinuous_28 |
| LTBP1 | tacatcgacacaggtcatct | LTBP1_exonContinuous_29 |
| LTBP1 | gttgttcaactcatcgcatt | LTBP1_exonContinuous_30 |
| LTBP1 | gtgttaatgcacttggcatt | LTBP1_exonContinuous_31 |
| LTBP1 | aggtcactgtctttctctaa | LTBP1_exonContinuous_32 |
| TGM2 | agatcacacctctctaagac | TGM2_exonContinuous_1 |
| TGM2 | cactgaaggtgagactgtct | TGM2_exonContinuous_2 |

| | | |
|---|---|---|
| TGM2 | ttagtggaaaacgggccttg | TGM2_exonContinuous_3 |
| TGM2 | gagtccaggtacacagcatc | TGM2_exonContinuous_4 |
| TGM2 | tggtagataaagccctgctg | TGM2_exonContinuous_5 |
| TGM2 | cttgatgaacttggccgagc | TGM2_exonContinuous_6 |
| TGM2 | gcccaaaattccaaggtatg | TGM2_exonContinuous_7 |
| TGM2 | tctaggatcccatcttcaaa | TGM2_exonContinuous_8 |
| TGM2 | catcgttgcagttgaccatg | TGM2_exonContinuous_9 |
| TGM2 | cagcactggccatacttgac | TGM2_exonContinuous_10 |
| TGM2 | gagttgtagttggtcacgac | TGM2_exonContinuous_11 |
| TGM2 | gatgagaaggttgctgttct | TGM2_exonContinuous_12 |
| TGM2 | ccaaactcattgcggaagta | TGM2_exonContinuous_13 |
| TGM2 | agcagtggaagttccagatc | TGM2_exonContinuous_14 |
| TGM2 | atggcacgaactggaactgg | TGM2_exonContinuous_15 |
| TGM2 | catcgtacttggtgctcagg | TGM2_exonContinuous_16 |
| TGM2 | attgacctccgcaaagacaa | TGM2_exonContinuous_17 |
| TGM2 | gatttgtgcacagacccatc | TGM2_exonContinuous_18 |
| TGM2 | caacgatcagggaacggttg | TGM2_exonContinuous_19 |
| TGM2 | ctcttagtgctgatcttcag | TGM2_exonContinuous_20 |
| TGM2 | ttgtaggtgtgggtgatatc | TGM2_exonContinuous_21 |
| TGM2 | ccagtttgttcaggtggttc | TGM2_exonContinuous_22 |
| TGM2 | tgttggtgatgtgggcaaag | TGM2_exonContinuous_23 |
| TGM2 | aagatcccattgtagctgac | TGM2_exonContinuous_24 |
| TGM2 | tgaggttgagcaggtacttg | TGM2_exonContinuous_25 |
| TGM2 | ttctcagagaaaggctccag | TGM2_exonContinuous_26 |
| TGM2 | gaggatgcaaagaggaacgc | TGM2_exonContinuous_27 |
| TGM2 | taaggcagtcacggtatttc | TGM2_exonContinuous_28 |
| TGM2 | accttgatgaggttggactc | TGM2_exonContinuous_29 |
| TGM2 | ataactggctccacgaggag | TGM2_exonContinuous_30 |
| TGM2 | caggtccattctcaccttaa | TGM2_exonContinuous_31 |
| TGM2 | ccaatgatgacattccggaa | TGM2_exonContinuous_32 |
| hUBC | atggtcttaccagtcagagt | hUBC_1 |
| hUBC | gacattctcgatggtgtcac | hUBC_2 |
| hUBC | gggatgccttccttatcttg | hUBC_3 |
| hUBC | atcttccagctgttttccag | hUBC_4 |
| hUBC | cagtgagtgtcttcacgaag | hUBC_5 |
| hUBC | tcctggatctttgctttgac | hUBC_6 |

| | | |
|---|---|---|
| hUBC | cagggtagactctttctgga | hUBC_7 |
| hUBC | cttcacgaagatctgcatcc | hUBC_8 |
| hUBC | tcttggatctttgccttgac | hUBC_9 |
| hUBC | cagtgagtgtcttcacgaag | hUBC_10 |
| hUBC | tgacgttctcgatagtgtca | hUBC_11 |
| hUBC | tccttgtcttggatctttgc | hUBC_12 |
| hUBC | cagggtagactctttctgga | hUBC_13 |
| hUBC | cttcacgaagatctgcatcc | hUBC_14 |
| hUBC | agagtgatggtcttaccagt | hUBC_15 |
| hUBC | tcttggatctttgccttgac | hUBC_16 |
| hUBC | cttcacgaagatctgcatcc | hUBC_17 |
| hUBC | agagtgatggtcttaccagt | hUBC_18 |
| hUBC | tcttggatctttgccttgac | hUBC_19 |
| hUBC | tgtttcccagcaaagatcaa | hUBC_20 |
| hUBC | cttcacgaagatctgcatcc | hUBC_21 |
| hUBC | agagtgatggtcttaccagt | hUBC_22 |
| hUBC | tcttggatctttgccttgac | hUBC_23 |
| hUBC | tgtttcccagcaaagatcaa | hUBC_24 |
| hUBC | cttcacgaagatctgcatcc | hUBC_25 |
| hUBC | agagtgatggtcttaccagt | hUBC_26 |
| hUBC | tcttggatctttgccttgac | hUBC_27 |
| hUBC | tgtttcccagcaaagatcaa | hUBC_28 |
| hUBC | gacattctcgatggtgtcac | hUBC_29 |
| hUBC | gggatgccttccttatcttg | hUBC_30 |
| hUBC | tgtttcccagcaaagatcaa | hUBC_31 |
| hUBC | agagtggactctttctggat | hUBC_32 |

**Table 4: Primers used to index ATAC sequencing libraries**

| **primer name** | **sequence** | **index** |
|---|---|---|
| N7001 | CAAGCAGAAGACGGCATACGAGATTCGCCTTAGTCTCGTGGGCTCGGAGATGTG | TAAGGCGA |
| N7002 | CAAGCAGAAGACGGCATACGAGATCTAGTACGGTCTCGTGGGCTCGGAGATGTG | CGTACTAG |
| N7003 | CAAGCAGAAGACGGCATACGAGATTTCTGCCTGTCTCGTGGGCTCGGAGATGTG | AGGCAGAA |
| N7004 | CAAGCAGAAGACGGCATACGAGATGCTCAGGAGTCTCGTGGGCTCGGAGATGTG | TCCTGAGC |
| N7005 | CAAGCAGAAGACGGCATACGAGATAGGAGTCCGTCTCGTGGGCTCGGAGATGTG | GGACTCCT |

| | | |
|---|---|---|
| N7006 | CAAGCAGAAGACGGCATACGAGATCATGCCTAGTCTCGTGGGCTCGGAGATGTG | TAGGCATG |
| N7007 | CAAGCAGAAGACGGCATACGAGATGTAGAGAGGTCTCGTGGGCTCGGAGATGTG | CTCTCTAC |
| N7008 | CAAGCAGAAGACGGCATACGAGATCCTCTCTGGTCTCGTGGGCTCGGAGATGTG | CAGAGAG G |
| N7009 | CAAGCAGAAGACGGCATACGAGATAGCGTAGCGTCTCGTGGGCTCGGAGATGTG | GCTACGCT |
| N7010 | CAAGCAGAAGACGGCATACGAGATCAGCCTCGGTCTCGTGGGCTCGGAGATGTG | CGAGGCT G |
| N7011 | CAAGCAGAAGACGGCATACGAGATTGCCTCTTGTCTCGTGGGCTCGGAGATGTG | AAGAGGC A |
| N7012 | CAAGCAGAAGACGGCATACGAGATTCCTCTACGTCTCGTGGGCTCGGAGATGTG | GTAGAGG A |
| N5001 | AATGATACGGCGACCACCGAGATCTACACTAGATCGCTCGTCGGCAGCGTCAGAT GTGTAT | TAGATCGC |
| N5002 | AATGATACGGCGACCACCGAGATCTACACCTCTCTATTCGTCGGCAGCGTCAGAT GTGTAT | CTCTCTAT |
| N5003 | AATGATACGGCGACCACCGAGATCTACACTATCCTCTTCGTCGGCAGCGTCAGAT GTGTAT | TATCCTCT |
| N5004 | AATGATACGGCGACCACCGAGATCTACACAGAGTAGATCGTCGGCAGCGTCAGA TGTGTAT | AGAGTAGA |
| N5005 | AATGATACGGCGACCACCGAGATCTACACGTAAGGAGTCGTCGGCAGCGTCAGA TGTGTAT | GTAAGGA G |
| N5006 | AATGATACGGCGACCACCGAGATCTACACACTGCATATCGTCGGCAGCGTCAGAT GTGTAT | ACTGCATA |
| N5007 | AATGATACGGCGACCACCGAGATCTACACAAGGAGTATCGTCGGCAGCGTCAGA TGTGTAT | AAGGAGTA |
| N5008 | AATGATACGGCGACCACCGAGATCTACACCTAAGCCTTCGTCGGCAGCGTCAGAT GTGTAT | CTAAGCCT |
| N5009 | AATGATACGGCGACCACCGAGATCTACACTGGAAATCTCGTCGGCAGCGTCAGAT GTGTAT | TGGAAATC |
| N5010 | AATGATACGGCGACCACCGAGATCTACACAACATGATTCGTCGGCAGCGTCAGAT GTGTAT | AACATGAT |
| N5011 | AATGATACGGCGACCACCGAGATCTACACTGATGAAATCGTCGGCAGCGTCAGAT GTGTAT | TGATGAAA |
| N5012 | AATGATACGGCGACCACCGAGATCTACACGTCGGACTTCGTCGGCAGCGTCAGA TGTGTAT | GTCGGAC T |
